# Regional reproducibility of immune expression programs in glioblastoma and implications for single-biopsy assessment

**DOI:** 10.64898/2026.10.07.757451

**Authors:** Corey Keith Goldman

**Affiliations:** NYU Langone Health / MedAlliance Medical Health Services, New York, NY, United States

**Keywords:** glioblastoma, tumor microenvironment, intratumoral heterogeneity, intraclass correlation coefficient, interferon, CD8, biomarkers, hepatocellular carcinoma

## Abstract

Tissue immune measurements can guide immunotherapy only if one biopsy represents the rest of the tumor. We quantified this in a secondary analysis of public data from 75 magnetic resonance imaging (MRI)-localized glioblastoma (GBM) biopsies from 19 patients. Fourteen transcriptomic modules scored supportive, inhibitory, and checkpoint programs. The intraclass correlation coefficient (ICC) gave the share of variance between patients rather than regions, before and after removing the component explained by a 12-gene leukocyte composite. An ICC of 1 indicates no variation in module scores among biopsies within each patient, whereas an ICC of 0 indicates that biopsies from the same patient are no more similar, on average, than biopsies from different patients. Two hepatocellular carcinoma (HCC) cohorts and a microdissected GBM atlas served as comparisons. CD8 had the lowest unadjusted ICC among abundance scores (0.42; 95% patient-bootstrap interval, 0.10 to 0.68; leukocytes, 0.65). Within patients, CD8 correlated with supportive and inhibitory modules alike (unadjusted mean r, 0.37 and 0.36) but not with major histocompatibility complex class I (MHC-I; r = 0.06). Leukocyte adjustment lowered the median module ICC from 0.56 to 0.34 but interferon (IFN) only from 0.74 to 0.70. IFN had the highest adjusted ICC and ranked first in 86% of patient-bootstrap resamples (post hoc). MHC-I was second (0.47) and not separable from IFN (difference 0.23; interval, −0.04 to 0.43); after purity residualization in 67 biopsies, IFN and MHC-I were similar (0.66 and 0.62). Median adjusted ICCs were 0.34 in bulk GBM, 0.62 and 0.74 in HCC, and 0.08 in microdissected GBM. In bulk GBM, only IFN, and MHC-I after purity residualization, had adjusted ICC point estimates above 0.5. The data support multiregion sampling with CD8, IFN, MHC-I, and checkpoint readouts from the same regions.

## Introduction

Immunotherapy depends on a sequence of events: release and presentation of tumor antigen, priming of T cells, trafficking of those cells into tumor tissue, and effector function that inhibitory signals do not block. The cancer-immunity cycle describes this sequence, and its 2023 update adds dendritic-cell support, checkpoint regulation, and the tumor microenvironment [1]. Any step can become rate-limiting [1], so treatment can strengthen one step while another still restricts the response. A tissue measurement that reports only immune-cell abundance cannot identify the limiting step.

Tissue measurements of programs, alongside abundance, have been associated with response to checkpoint blockade. Pretreatment CD8-positive cell localization and expression of programmed cell death protein 1 (PD-1) and its ligand (PD-L1) were associated with response to pembrolizumab in melanoma [2]. An interferon-gamma-related expression profile was associated with response to PD-1 blockade across several cancer cohorts [3]. Stromal expression of transforming growth factor beta (TGF-β) and T-cell exclusion were associated with resistance to atezolizumab in urothelial cancer [4]. Clinical trials have incorporated tissue pharmacodynamic measurements and matched biomarkers incompletely [5,6], and the 2025 Society for Immunotherapy of Cancer (SITC) consensus called for more consistent biomarker collection and reporting [7].

Immune composition differs between regions of the same tumor in glioblastoma (GBM), lung adenocarcinoma, and melanoma [8–12]. In GBM, spatial and single-cell studies place distinct myeloid programs in hypoxic, vascular, and inflammatory niches [13,14], and the perinecrotic niche shows marked immunosuppression [15]. Ravi and colleagues found HMOX1-positive myeloid cells beside dysfunctional T cells in mesenchymal tumor regions and implicated myeloid interleukin (IL)-10 in T-cell dysfunction [16]. Regional differences therefore involve the programs that accompany immune cells as well as their abundance.

Spatial maps show where programs occur. A treatment decision needs a different quantity: how closely one biopsy reproduces a patient’s measurement relative to differences between patients. In hepatocellular carcinoma (HCC), Shen and colleagues evaluated this for single-region immune measurements by immunohistochemistry and RNA sequencing [17], and Zhang and colleagues developed a signature to reduce regional misclassification [18]. For GBM, we did not find a study that estimated how reproducibly magnetic resonance imaging (MRI)-localized biopsies capture a fixed panel of immune programs, or that separated that reproducibility from variation in leukocyte abundance. A 2026 reporting framework for spatial omics in high-grade glioma identifies sampling bias as one driver of divergent findings [19].

We analyzed public multiregion transcriptomic data from 75 MRI-localized GBM biopsies (19 patients) and, for comparison, a microdissected GBM atlas and two HCC cohorts. Fourteen predefined expression modules (nine supportive, four inhibitory, and one checkpoint module) scored programs of the cancer-immunity cycle. The intraclass correlation coefficient (ICC) estimated the share of variance lying between patients, before and after removing the component explained by a leukocyte composite. An ICC of 1 indicates no variation in module scores among biopsies within each patient, whereas an ICC of 0 indicates that biopsies from the same patient are, on average, no more similar than biopsies from different patients. Higher ICC therefore supports using a single biopsy to characterize that module; lower ICC reduces confidence that one biopsy represents the patient’s tumor. In GBM, interferon (IFN)-associated expression had the highest leukocyte-adjusted ICC (0.70), followed by major histocompatibility complex class I (MHC-I; 0.47), and the data did not separate IFN from MHC-I. The other 12 modules had adjusted ICCs of 0.42 or lower. Within patients, CD8 abundance correlated with supportive and inhibitory programs alike.

## Results

### Four sampling designs separated variation between patients from variation between regions

The primary analysis used 75 nonnecrotic bulk biopsies from 19 patients with MRI-localized GBM [20]. The comparison cohorts were 122 microdissected samples from 10 tumors in the Ivy Glioblastoma Atlas (GBM-IVY) [21] and two multiregion HCC cohorts, with 41 regions from 12 patients [17] and 75 regions from 14 patients [22] (Table 1). Every cohort was scored on 14 modules (Table 2; gene membership in Supplementary Table S1). The four inhibitory modules describe programs that restrict immune-cell access or function: angiogenic, stromal/TGF-β, myeloid, and regulatory T cell (Treg).

**Table 1.** Cohorts and sampling questions.

| Cohort label | Regional analysis | Sampling design and purpose |
| --- | --- | --- |
| GBM | 19 patients, 75 regions | MRI-localized bulk biopsies. Tests how representative a regional biopsy is of the patient's immune profile. Descriptive population: 23 patients, 79 regions. |
| GBM-IVY | 10 tumors, 122 samples | Five laser-capture anatomical structures. Tests whether microscopic environments differ within the same tumor. |
| HCC cohort 1 | 12 patients, 41 regions | Multiregion bulk samples. Independent solid-tumor comparison for regional consistency. |
| HCC cohort 2 | 14 patients, 75 regions | Multiregion bulk samples with spatial coordinates. Second independent HCC comparison. |

The GBM primary and regional-analysis populations overlap and must not be added. GBM-IVY samples have a different anatomical sampling design. Repository identifiers and processing details are given in Methods.

**Table 2.** Why the 14 immune modules were examined.

| Module | Relationship to the extended cancer-immunity cycle | Biological question retained by measuring it separately |
| --- | --- | --- |
| IFN/STING | Innate sensing and interferon-associated activation | Is an activation-associated transcriptional program present across sampled regions? |
| MHC-I presentation | Antigen processing and display for T-cell recognition | Does presentation machinery accompany immune activation? |
| cDC1/APC | Cross-presentation and continued dendritic-cell support | Is the dendritic-cell-associated program preserved where effector cells are present? |
| Th1/IL-12 | Type 1 polarization and support of effector responses | Does a type 1 support program accompany presentation and cytotoxic expression? |
| CD8/cytotoxic | Lymphocyte-associated expression and killing machinery | Are effector-associated transcripts present beyond the CD8 lineage score alone? |
| NK/IL-15 | NK-associated effector biology and IL-15-related support | Does a complementary effector-support program accompany the T-cell profile? |
| Memory/IL-7 | Naive-memory and survival-associated features | Are repertoire and persistence-associated transcripts relatively preserved? |
| Tfh/IL-21 | Helper and persistence-associated features | Is helper-associated support present alongside activation and effectors? |
| Trafficking/adhesion | Recruitment, adhesion, and access to tumor tissue | Do immune cells have a corresponding recruitment and adhesion program? |
| Adaptive checkpoint | Inhibitory receptors and ligands accompanying immune engagement | Is checkpoint-associated expression present where CD8 and supportive programs are expressed? |
| Angiogenic | Vascular programs that may affect immune entry | Could vascular organization constitute a distinct access barrier? |
| Stromal/TGF-β | Matrix, stromal, and TGF-β-associated programs | Could tissue organization or stromal regulation limit immune access or function? |
| Myeloid | Tumor-associated macrophage and myeloid-derived suppressor cell programs | Does myeloid-associated expression coexist with activation and lymphocytes? |
| Treg | Regulatory T-cell-associated programs | Is regulatory-cell-associated expression present across sampled regions or concentrated in particular regions? |

Each module was analyzed on two scales: the unadjusted bulk score, and the residual score after removing its fitted relationship with a 12-gene leukocyte composite. Leukocyte, CD3, and CD8 abundance scores were analyzed separately.

### CD8 abundance had the lowest ICC of the three abundance scores in GBM

Among the three unadjusted abundance scores, CD8 had the lowest GBM ICC: 0.42 (95% patient-bootstrap interval, 0.10 to 0.68). The ICCs were 0.55 (0.33 to 0.72) for CD3 and 0.65 (0.49 to 0.77) for leukocytes (Figure 1A). The paired leukocyte-minus-CD8 difference was 0.23 (−0.04 to 0.53), so the ordering is uncertain (Supplementary Table S7). The CD8 score uses two genes and the leukocyte score twelve. Scores built from each of the 66 leukocyte gene pairs had a median ICC of 0.60 (2.5th to 97.5th percentile, 0.42 to 0.70). Of these pairs, 97% exceeded the CD8 ICC of 0.42 (Supplementary Table S10F). Gene count therefore does not explain the lower CD8 point estimate.

**Figure 1.**
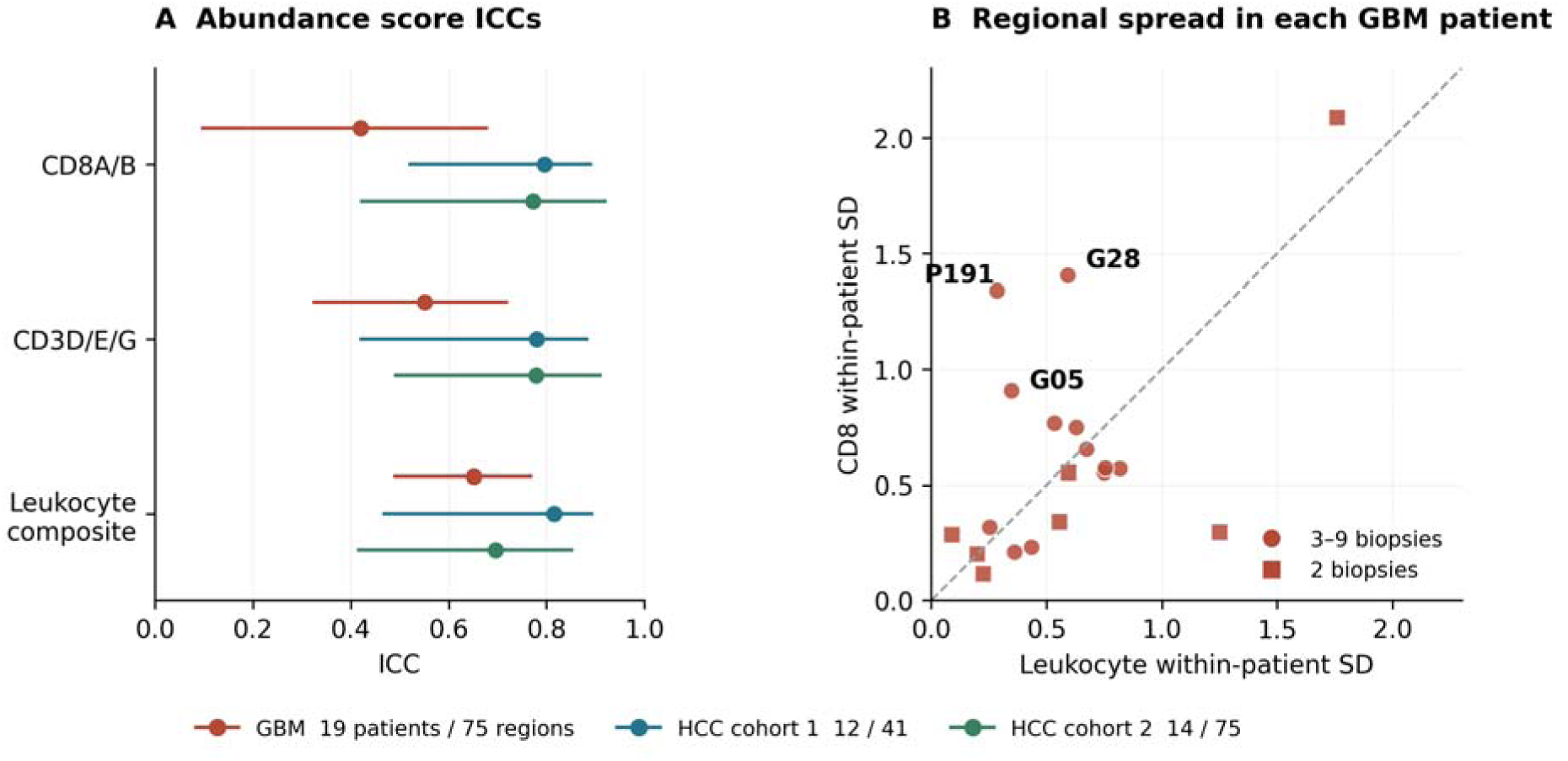
CD8 abundance had the lowest GBM ICC among abundance scores, with regional spread concentrated in a few patients. (A) ICCs for CD8, CD3, and leukocyte scores (unadjusted), with pointwise 95% intervals from 5,000 patient bootstraps. GBM includes 19 patients and 75 regions; HCC cohorts 1 and 2 include 12 patients and 41 regions and 14 patients and 75 regions. (B) Each GBM patient’s CD8 SD plotted against leukocyte SD; the dashed line marks equal SDs. Squares denote patients with two biopsies and circles three to nine. G28, P191, and G05 show greater CD8 than leukocyte spread. The paired leukocyte-minus-CD8 ICC interval includes zero, so the ordering is uncertain (Supplementary Table S7). Scores were standardized within their source references.

Both HCC cohorts had higher CD8 ICCs: 0.80 (0.52 to 0.89) and 0.77 (0.42 to 0.92). The HCC-minus-GBM differences were 0.38 (0.02 to 0.70) for cohort 1 and 0.35 (−0.09 to 0.70) for cohort 2, and only the first interval excluded zero. With exactly two regions sampled per patient, median CD8 ICCs were 0.46 in GBM and 0.78 and 0.72 in HCC cohorts 1 and 2.

In GBM, the lower CD8 ICC reflected less between-patient variance than the leukocyte composite (0.51 versus 0.80) and more within-patient variance (0.70 versus 0.43), both on the unadjusted scale. Regional spread differed among patients (Figure 1B). Patients G28 and P191 contributed 35% and 14% of the CD8 within-patient sum of squares, from eight and four biopsies. Omitting G28 raised the CD8 ICC from 0.42 to 0.55. Supplementary Tables S2 and S8 give abundance summaries and per-patient standard deviations (SDs).

### CD8 abundance tracked supportive and inhibitory programs alike and carried little information about MHC-I

To ask whether a CD8-rich region also expresses supportive programs, we correlated each biopsy’s deviation from its patient’s mean CD8 score with its deviations in each module score (Figure 2). Every patient received equal weight. The correlation between CD8 and MHC-I expression was 0.06 (95% interval, −0.14 to 0.28) and stayed near zero when any single patient was omitted. MHC-I tracked the leukocyte composite (r = 0.42) and CD3 (r = 0.28) more closely than CD8. The unadjusted within-patient variance of MHC-I (0.52 squared standardized units) was not negligible, so the weak CD8 association did not arise from an invariant MHC-I score.

**Figure 2.**
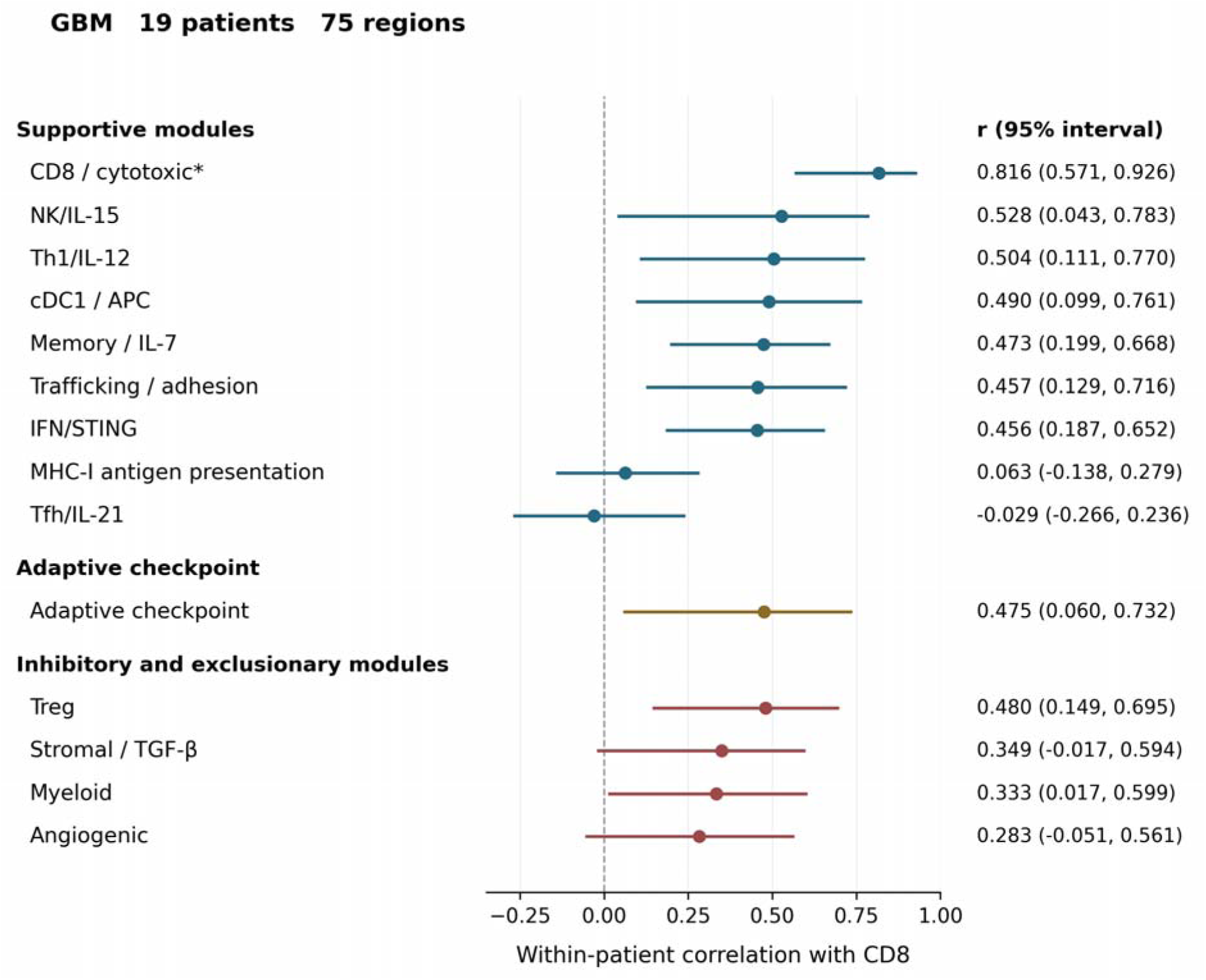
CD8 expression correlated with supportive and inhibitory programs alike in GBM, and only weakly with MHC-I. Points show correlations between the CD8A/B expression score and all 14 unadjusted module scores in 75 GBM biopsies from 19 patients. Both variables were centered within patient, and each patient received equal total weight. Bars are pointwise 95% intervals from 5,000 patient bootstraps. Modules are grouped by category (supportive, inhibitory, checkpoint) and ordered from highest to lowest correlation within each category. MHC-I expression showed little regional association with CD8, whereas checkpoint, Treg, and myeloid scores had positive associations. The mean correlation was 0.37 across the eight supportive modules that share no genes with CD8 and 0.36 across the four inhibitory modules. The ranking is descriptive and does not establish statistically distinct positions. The asterisk marks the CD8A/B genes shared with the mixed cytotoxic module, so its high correlation depends partly on that overlap. These are RNA expression measurements and not direct cell counts or functional assays. Full three-cohort results are in Supplementary Figure S2 and Table S4.

The conventional type 1 dendritic-cell (cDC1) module correlated with CD8 (r = 0.49; 0.10 to 0.76). CD8 correlations were also positive for the checkpoint, Treg, and myeloid modules. Stromal/TGF-β and angiogenic correlations were positive with intervals that included zero. The T follicular helper cell (Tfh)/IL-21 module, scored from five of nine defined genes, showed another near-zero association.

The CD8/cytotoxic module had the highest CD8 correlation (r = 0.82), partly because both scores contain CD8A and CD8B. Removing six lymphocyte-lineage genes (Methods) lowered the correlation to 0.64 while retaining granzyme, perforin, and other effector transcripts. The remaining correlation links lymphocyte abundance to effector-gene expression and does not measure cytotoxic activity.

Thirteen of 14 unadjusted module correlations with CD8 were positive, spanning supportive and inhibitory programs. In an exploratory contrast, the mean unadjusted CD8 correlation was 0.37 across the eight supportive modules that share no genes with CD8 and 0.36 across the four inhibitory modules. The supportive-minus-inhibitory difference was 0.01 (95% patient-bootstrap interval, −0.12 to 0.16). After leukocyte adjustment the means were 0.115 and 0.190 (difference, −0.075; −0.19 to 0.07). Leukocyte adjustment weakened several of the individual correlations. The IFN, checkpoint, and trafficking correlations with CD8 fell from 0.46, 0.48, and 0.46 to 0.09, 0.18, and 0.08. The unadjusted correlations of the same three modules with the leukocyte composite were 0.61, 0.64, and 0.85 (Supplementary Figure S3). Much of the CD8 covariation with these modules therefore accompanied broad leukocyte expression, and CD8 abundance did not separate supportive from inhibitory programs. On the unadjusted scale, the GBM tests in the restricted 42-test family gave q = 0.055 for IFN and trafficking and q = 0.161 for checkpoint (Supplementary Table S5).

In HCC, unadjusted checkpoint correlations with CD8 were 0.94 and 0.80 and trafficking correlations were 0.87 and 0.78 in cohorts 1 and 2. All four met q < 0.012 across the 42-test family, whereas no adjusted association in that family met q < 0.05 (Supplementary Table S5). Supplementary Figure S2 and Table S4 show all 14 unadjusted modules against both abundance scores.

### IFN-associated expression kept its ICC after leukocyte adjustment, whereas cDC1, trafficking, and myeloid programs did not

Leukocyte adjustment lowered the median GBM module ICC from 0.56 to 0.34 (Figure 3A), but IFN changed only from 0.74 to 0.70 (adjusted 95% interval, 0.44 to 0.80). The cDC1, trafficking, and myeloid modules, which track leukocyte expression closely, fell from 0.60, 0.64, and 0.59 to 0.22, 0.18, and 0.29. The unadjusted ICC describes the bulk module score. The adjusted ICC describes the residual score after removing its fitted relationship with the leukocyte composite.

**Figure 3.**
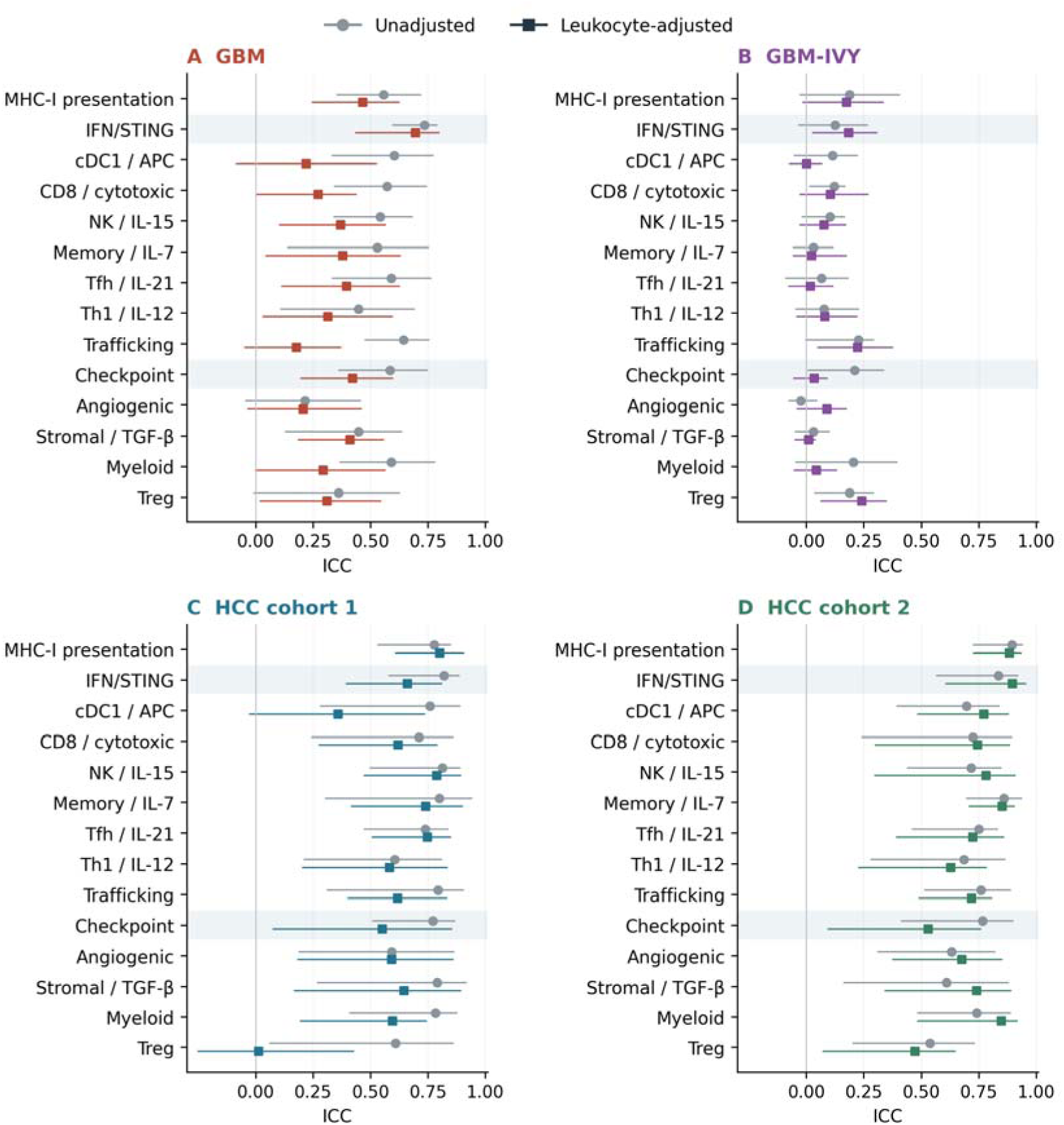
Module ICCs differ by immune program and sampling scale. Unadjusted and leukocyte-adjusted ICCs for all 14 modules in (A) GBM, (B) GBM-IVY, (C) HCC cohort 1, and (D) HCC cohort 2. Gray circles show unadjusted values, colored squares show leukocyte-adjusted values, and lines show pointwise 95% patient-bootstrap intervals. IFN and checkpoint rows are shaded. IFN has the highest adjusted GBM point estimate; all 13 IFN contrasts are in Supplementary Table S7. GBM-IVY samples microscopic structures, whereas the other cohorts use bulk regions. Exact estimates and variance components are in Supplementary Table S3.

IFN ranked first among the 14 adjusted modules in 86% of patient-bootstrap resamples and among the top three in 96%; on the unadjusted scale it ranked first in 72%. Among the 13 IFN contrasts, simultaneous intervals excluded zero for the adjusted CD8/cytotoxic, trafficking, and angiogenic modules but not for cDC1 or myeloid (Supplementary Table S7). The IFN-minus-MHC-I contrast was 0.23 (pointwise 95% interval, −0.04 to 0.43), so the data do not separate IFN from MHC-I. The ranks are descriptive frequencies from a post hoc comparison, not probabilities that IFN is best.

The IFN ranking persisted among the 12 patients with at least three biopsies (61 regions). The median adjusted ICC was 0.37, IFN remained first at 0.71, and the unadjusted CD8 and leukocyte ICCs were 0.44 and 0.66.

IFN covaried with other programs within patients. Adjusted IFN and MHC-I scores correlated across patient means (r = 0.76) and within patients (r = 0.55; 95% interval, 0.35 to 0.75). The within-patient correlation between adjusted IFN and checkpoint scores was 0.60 (0.33 to 0.78), compared with 0.76 before adjustment. The IFN (22 genes), MHC-I (13), and checkpoint (8) modules share no genes (Supplementary Table S1), so these correlations do not arise from common transcripts. The modules are nevertheless not independent, because each contains interferon-inducible genes (Methods). Removing the interferon-inducible genes CD274 and IDO1 from the eight-gene checkpoint module lowered its adjusted ICC from 0.42 to 0.26, and also removing PDCD1LG2 lowered it to 0.15. Without the three genes, the within-patient correlation between adjusted IFN and checkpoint scores fell from 0.60 to 0.46 (95% interval, 0.17 to 0.66) and stayed positive (Supplementary Table S10D). Interferon-inducible genes therefore account for part, but not all, of the IFN-checkpoint correlation.

### Controls for tumor purity, module size, and patient-level determinants left IFN first but did not separate it from MHC-I

Hu and colleagues estimated tumor purity from whole-exome sequencing (WES) [20]. Among 72 of the 79 descriptive-set biopsies (22 patients) with a purity estimate, adjusted MHC-I correlated positively with purity (Spearman rho = 0.40; Benjamini-Hochberg q = 0.012). The memory/IL-7 and type 1 helper T cell (Th1)/IL-12 modules correlated negatively (rho = −0.37 and −0.43; q = 0.036 and 0.013). IFN did not (rho = 0.09; q = 0.70) (Supplementary Table S9). In the 67 multiregion biopsies from 17 patients, residualizing the adjusted scores on purity raised the MHC-I ICC from 0.46 to 0.62 and the myeloid ICC from 0.26 to 0.39. The MHC-I change was 0.15 (95% interval, −0.06 to 0.32). The IFN ICC changed from 0.68 to 0.66. The IFN-minus-MHC-I difference fell from 0.21 (95% patient-bootstrap interval, −0.11 to 0.43) to 0.04 (−0.25 to 0.30), and IFN was highest of the 14 modules in 80% of resamples before residualization and 55% after. IFN and MHC-I were not separable before or after residualization, and no other module exceeded 0.42 after residualization.

IFN is the largest module, with 22 scored genes against 5 to 15 for the others. When every module was reduced to five randomly drawn genes (1,000 draws), the median adjusted ICC was 0.65 for IFN (2.5th to 97.5th percentile, 0.40 to 0.76) and 0.47 for MHC-I. The other modules ranged from 0.21 to 0.40 (Supplementary Table S10A). In 93% of the five-gene IFN subsets, the ICC exceeded the full-gene MHC-I ICC. The leukocyte composite was a reliable covariate. Random six-versus-six splits of its 12 genes correlated at r = 0.90 in the 141-biopsy reference (Spearman-Brown reliability, 0.95) and at r = 0.84 within patients (reliability, 0.91; Supplementary Table S10B). Scaling the fitted slope from 0 to 1.5 times its reference value changed the IFN ICC only from 0.74 to 0.65. Adding CD3 as a second covariate gave an IFN ICC of 0.69 against 0.43 for the next module (Supplementary Table S10C).

Two patient-level factors, 9p21 loss and glucocorticoid exposure, could create between-patient variance that does not reflect regional biology. Homozygous deletions of 9p21.3 eliminate CDKN2A and, in about half of cases, also a type I interferon gene cluster [23]. In these tumors, CDKN2A loss, a marker of 9p21 deletion, was frequent (19 of 22 patients; homozygous loss in 15). Patients with homozygous loss had higher, not lower, mean adjusted IFN scores than the other seven (0.39 versus −0.18; Mann-Whitney p = 0.27). The same held for any loss (0.34 in 19 patients versus −0.62 in three; p = 0.13). Adjusting the IFN score for CDKN2A and IFNB1 (the interferon-beta gene) copy number changed its ICC from 0.68 to 0.69 in the 67 multiregion biopsies with copy-number calls (Supplementary Table S10E). For glucocorticoid exposure, dexamethasone dose and duration were not recorded. A seven-gene glucocorticoid-response score, an unvalidated proxy, had its own patient-level component (unadjusted ICC 0.44) and correlated with the leukocyte composite across patient means (rho 0.52). Patient means of the score did not track IFN (rho = 0.18; p = 0.47), and residualizing IFN on the score left its ICC at 0.71 (Supplementary Table S10E).

### Microdissected GBM structures had lower ICCs than bulk biopsies

GBM-IVY had a median adjusted module ICC of 0.08 across its five canonical anatomical structures, with adjusted IFN and checkpoint ICCs of 0.18 and 0.03 (Figure 3B). The median pairwise correlation between 14-module adjusted profiles, which asks whether the same modules rank high or low in different samples from one tumor, was 0.07, compared with 0.31 in bulk GBM. In patient-centered anatomical summaries, microvascular proliferation had the highest IFN, cDC1, cytotoxic, and angiogenic scores. Viable pseudopalisading tissue around necrosis had the highest checkpoint and myeloid scores and the lowest IFN and cytotoxic scores. Laser-capture microdissection separates cellular environments that a bulk biopsy averages, so the two designs sample different tissue volumes.

Among 11 bulk GBM patients with both contrast-enhancing (CE) and nonenhancing (NE) samples, no adjusted module difference passed correction across the 14 programs. NE samples nevertheless had lower checkpoint and trafficking scores in 10 of 11 patients, with mean differences of −0.82 and −0.56 standardized units (Benjamini-Hochberg q = 0.123 and 0.146). Conditioning the ICC on region type left IFN first (0.71 by the method of moments, 0.68 by restricted maximum likelihood). The median module ICC was 0.36 (Supplementary Table S11).

### HCC cohorts had higher ICCs overall, with exceptions among inhibitory programs

Median adjusted module ICCs were 0.62 in HCC cohort 1 and 0.74 in cohort 2, compared with 0.34 in GBM (Figure 3C, D). Twelve of 14 modules had higher adjusted point estimates in both HCC cohorts. Adjusted estimates were 0.62 and 0.72 for trafficking, 0.36 and 0.77 for cDC1, and 0.60 and 0.85 for myeloid. IFN estimates were 0.66 and 0.90, so GBM (0.70) fell between the two HCC cohorts for this module.

Checkpoint and Treg did not follow the overall HCC ranking. Unadjusted checkpoint ICCs were 0.77 in both HCC cohorts, but adjusted estimates were 0.55 (95% interval, 0.08 to 0.85) and 0.53 (0.10 to 0.76). HCC cohort 1 had an adjusted Treg ICC of 0.01 against 0.61 unadjusted. A higher median ICC therefore did not extend to every inhibitory program.

Agreement in module ordering was also higher in HCC. Median pairwise correlations between 14-module adjusted profiles were 0.69 in both HCC cohorts and 0.31 in GBM; with exactly two regions sampled per patient they were 0.73 and 0.76 in HCC and 0.34 in GBM. The HCC-GBM comparisons are conditional on anatomical sampling, tissue volume, processing, and the between-patient variance of each cohort. Matching the number of biopsies does not remove those differences, and regional SDs on separately standardized scales are not absolute expression differences between cancer types.

### Patients differed in how well one biopsy represented the rest of the tumor

To see how cohort-level values appear in single tumors, we compared the 19 GBM patients on two summaries of regional score variation. Profile correlation is the median pairwise Spearman correlation between the 14 adjusted module scores across a patient’s biopsies, and is high when the same modules rank high or low in every region. Centroid root mean square error (RMSE) is the median distance of each biopsy from the patient’s median profile, and is low when regional scores are close. Neither is an individual-patient ICC. Patient G35 had the highest profile correlation (0.73) and the lowest centroid RMSE (0.29), and patient P191 had the lowest correlation (−0.18) and the highest RMSE (1.26) (Supplementary Figure S4 and Table S8). The same extremes held among the 12 patients with at least three biopsies.

In G35, the adjusted cDC1, checkpoint, and Treg scores exceeded the reference mean in all three regions, and no other patient among the 19 met this criterion. The reference mean is the mean of the 141-biopsy MRI reference used for standardization. Leukocyte and CD8 scores also exceeded the reference in all three G35 biopsies. A biopsy from any G35 region would therefore have shown dendritic-cell-associated expression coexisting with inhibitory programs. Adjusted IFN scores in G35 nevertheless ranged from −0.47 to 0.88, so a high cohort-level IFN ICC did not imply consistent IFN expression in every patient.

In P191, CD8, checkpoint, and stromal/TGF-β scores differed substantially between regions, whereas leukocyte scores varied less: leukocyte scores ranged from 0.46 to 1.12 and CD8 scores from −0.96 to 1.90. The two CE samples had higher adjusted checkpoint and stromal/TGF-β scores than the two NE samples, and cytotoxic and Th1 scores also differed. A single biopsy from P191 could therefore have missed a region with higher CD8 expression accompanied by checkpoint and stromal expression. The checkpoint direction agrees with the paired CE-versus-NE summary, while the magnitude and full profile are specific to this patient.

Similar mean expression did not imply similar regional consistency. Mean adjusted memory/IL-7 scores were 1.45 in G35 and 1.47 in P191, but the regional ranges were 1.25 to 1.69 and 0.40 to 2.48. The patients were not selected for matching averages across modules; this module shows that expression magnitude and regional spread are separate properties. Figures 4 and 5 place both patients on common axes before and after leukocyte adjustment, Supplementary Figure S1 shows the same values as radar plots, and Supplementary Table S6 lists every biopsy score.

**Figure 4.**
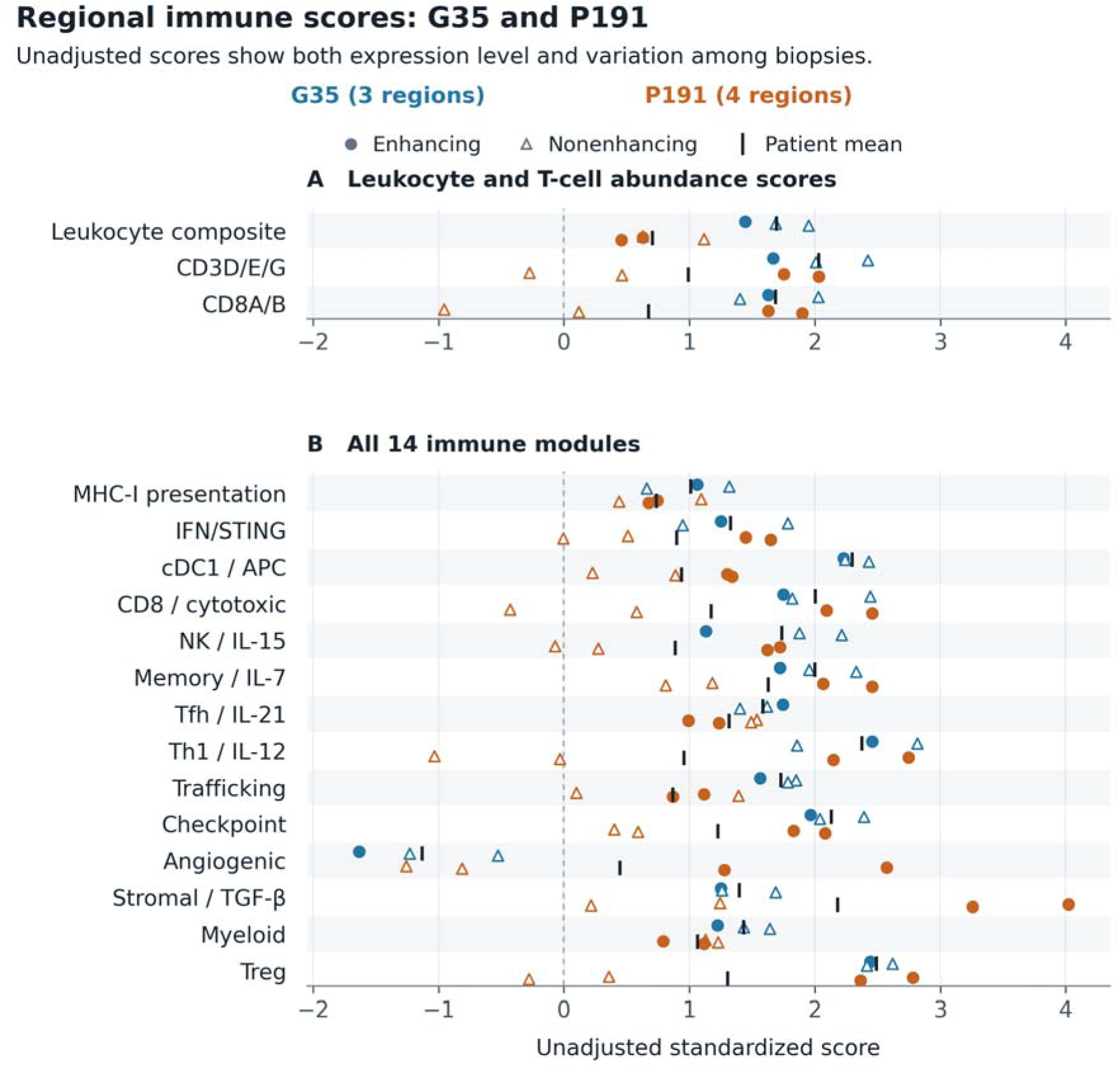
G35 and P191 differ in regional variation before leukocyte adjustment. (A) Leukocyte, CD3, and CD8 abundance scores. (B) All 14 unadjusted immune-module scores. Each point is one biopsy; G35 (blue, three biopsies) is shown above P191 (orange, four biopsies) in each row. Filled circles identify CE regions and open triangles NE regions. Black vertical ticks mark each patient’s arithmetic mean. The horizontal position gives the score, and the spread of a patient’s points shows regional variation. The dashed vertical line marks the reference mean and not a clinical cutoff. P191 varies much more in CD8 and several modules than in the leukocyte composite. All biopsy values are in Supplementary Table S6.

**Figure 5.**
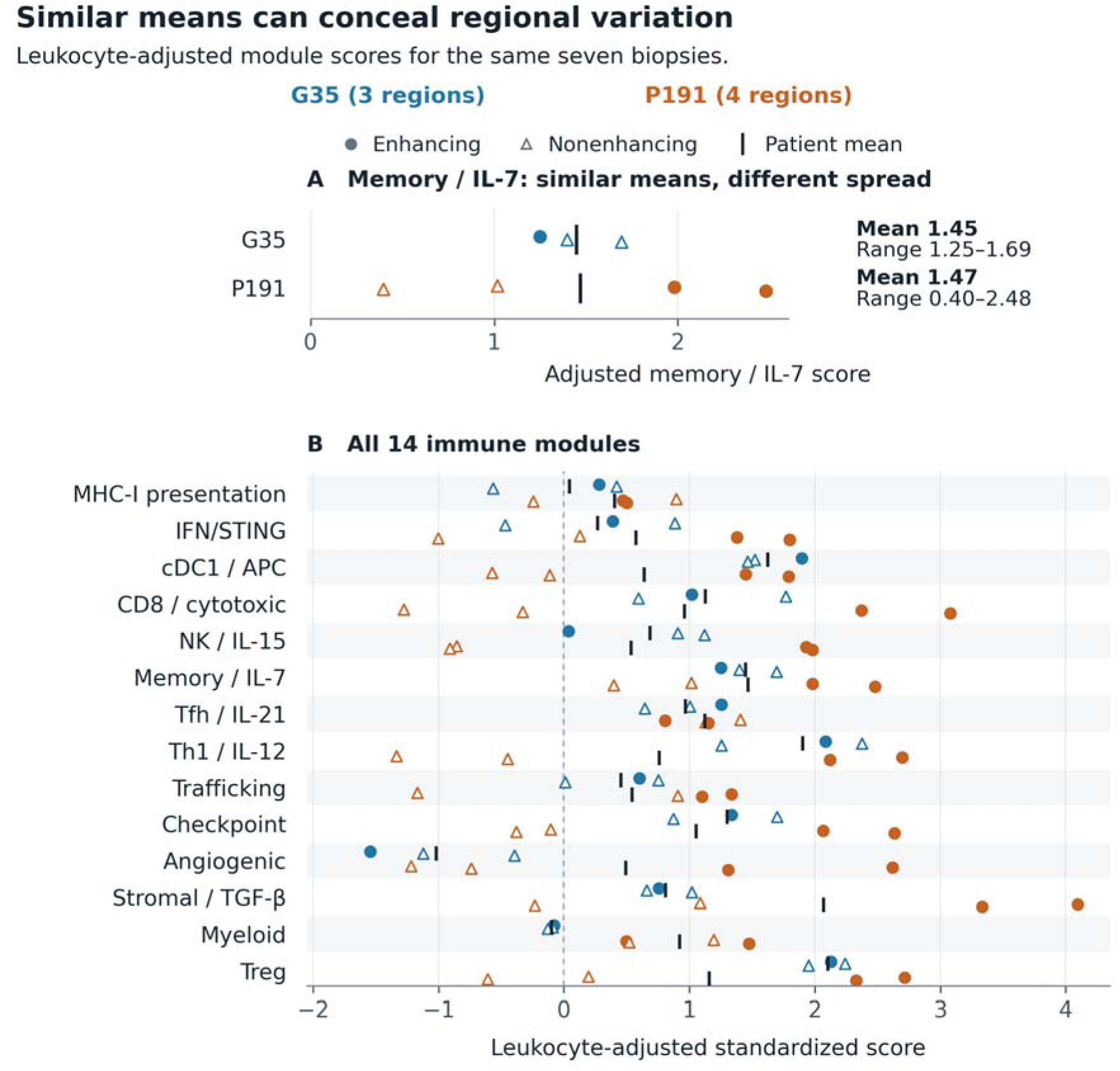
Similar mean module scores can conceal different regional variation. (A) Mean leukocyte-adjusted memory/IL-7 scores are similar in G35 and P191 (1.45 and 1.47), but the observed regional ranges differ (1.25 to 1.69 and 0.40 to 2.48). (B) All 14 adjusted modules, including memory/IL-7. Each point is one biopsy; blue identifies G35 and orange P191, with G35 above P191 in each row. Filled circles identify CE regions and open triangles NE regions. Black ticks mark arithmetic means over the sampled biopsies. The dashed line in (B) marks the reference mean and not a clinical cutoff. Unadjusted scores for the same biopsies are in Figure 4, and exact values are in Supplementary Table S6. The patients were not matched on overall module expression.

### Additional biopsies improved agreement with other regions of the same tumor

Six patients had at least five eligible biopsies. For each, we compared the mean profile of one to three selected biopsies with the mean profile of two other biopsies from the same patient, across all admissible splits. Profile correlation improved at each step in all six patients, with cohort medians of 0.28, 0.32, and 0.42 for one, two, and three selected biopsies. Profile errors and a threshold-based secondary comparison are in the Supplementary Methods. Quality-control exclusions changed the median adjusted GBM ICC by about 0.02, and leave-one-patient-out adjusted IFN ICCs ranged from 0.64 to 0.75.

A projection from the ICC with the Spearman-Brown formula gives the number of regions whose mean would reach a reliability, ICC(1,k), of 0.80. The point estimates are 2 regions for adjusted IFN, 5 for MHC-I, 6 for checkpoint, 8 for the median module, 15 for cDC1, and 19 for trafficking. For unadjusted leukocyte, CD3, and CD8 abundance they are 3, 4, and 6 (Supplementary Table S12). The projection treats regions as exchangeable draws from each patient. MRI-targeted sampling of CE and NE regions is not exchangeable, and intervals for the projected numbers are wide. The figures therefore rank programs by the sampling each needs and do not state a required number of biopsies.

## Discussion

In GBM, one biopsy reproduces some immune-expression programs of a patient more faithfully than others. IFN-associated expression had the highest leukocyte-adjusted ICC (0.70; 95% interval, 0.44 to 0.80), and MHC-I was second (0.47). The interval for the IFN-minus-MHC-I difference included zero, and after residualizing on tumor purity in 67 biopsies, IFN and MHC-I were similar (0.66 and 0.62). The remaining modules had adjusted ICCs of 0.18 to 0.42. The cDC1, trafficking, and myeloid programs lost more than half of their ICC once leukocyte abundance was removed. On the adjusted scale, IFN was the only GBM module whose 95% interval reached the good band under conventional reliability bands (above 0.75), and its lower limit lay in the poor band [24]. Within patients, unadjusted CD8 correlations averaged 0.37 with supportive and 0.36 with inhibitory modules. Median adjusted ICCs were higher in both HCC cohorts and lower in microdissected GBM than in bulk GBM. What one biopsy reveals about the rest of the tumor therefore differed with the program measured, the scale on which it was scored, and the sampling design.

The weak regional association between CD8 and MHC-I supports measuring antigen-presentation machinery separately from effector-cell abundance. MHC-I covaried with leukocyte expression and, more weakly, with CD3, yet showed little association with CD8. Bulk RNA cannot assign MHC-I transcripts to tumor, myeloid, or lymphoid cells, so this finding concerns composition and expression and does not demonstrate impaired antigen presentation. Adjusted MHC-I also tracked WES-inferred tumor purity across biopsies (rho = 0.40), and residualizing on purity raised its adjusted ICC from 0.46 to 0.62. This pattern is consistent with regional differences in tumor-cell content contributing to the within-patient variance of MHC-I. The interval for the change included zero, so that contribution remains a hypothesis. The positive CD8 correlations with supportive and inhibitory modules alike show that CD8 abundance did not indicate which accompanying programs were present. Spatial mapping and ex vivo experiments implicating myeloid interleukin-10 in local T-cell dysfunction show one route by which T-cell presence and T-cell function can diverge [16].

The IFN module retained its ICC after leukocyte adjustment and under alternative adjustment strengths, which indicates a stable patient-level component of IFN-associated expression beyond leukocyte abundance. Module size, noise in the leukocyte composite, and tumor purity did not account for that component. Two patient-level candidates were tested less definitively: CDKN2A loss, which was absent in only three patients, and glucocorticoid exposure, which was tested through an unvalidated expression proxy. Co-deletion of the interferon gene cluster within 9p21.3 disrupted type I interferon signaling and promoted immune evasion in a syngeneic mouse model of pancreatic cancer [23]. Here, CDKN2A loss, a marker of 9p21 deletion, was frequent and did not accompany lower IFN-associated expression. Dexamethasone dose and duration were not recorded, and the glucocorticoid-response score tracked the leukocyte composite, so dexamethasone and other patient-level exposures remain candidate determinants of the between-patient component. We propose, as a hypothesis for testing, that diffusible interferon signaling coordinates expression across regions while local vessels, stroma, and immune cells shape the other programs. The expression data do not test that mechanism. The IFN correlation with MHC-I did not arise from shared genes and could reflect coordinated interferon-responsive transcription in tumor and nontumor cells. Leukocyte adjustment cannot isolate tumor-cell-intrinsic signaling, and the IFN/STING (stimulator of interferon genes) module is not an assay of STING activation and shares STAT1 and CXCL9 with the IFN-γ-related pembrolizumab signature [3].

Within a tumor, regions with higher adjusted IFN expression than that patient’s average also had higher adjusted checkpoint expression (within-patient r = 0.60). Interferon-associated expression, which can accompany antitumor immunity [3], therefore coexists with counter-regulatory receptors and ligands in the same regions. Part of this coexistence reflects shared regulation, because CD274, IDO1, and PDCD1LG2 are interferon-inducible genes of an interferon-gamma-related profile [3]. Without them, the adjusted checkpoint ICC fell from 0.42 to 0.15 and the within-patient IFN-checkpoint correlation fell from 0.60 to 0.46. The coexistence of interferon-associated and checkpoint expression resembles adaptive immune resistance in melanoma [2] and the interferon-dependent multigenic resistance program described in experimental models [25]. In a phase 1 trial, intratumoral MEDI9197, a toll-like receptor 7/8 agonist, increased CD8-positive and PD-L1-positive cells and induced type 1 and 2 interferons without objective responses [26]. The trial pattern is consistent with, but does not establish, interferon induction accompanied by checkpoint upregulation, and our data do not show that treatment induces checkpoint expression. Stromal TGF-β has been linked to T-cell exclusion in urothelial cancer [4], and effector, Treg, vascular, and myeloid features to response to atezolizumab plus bevacizumab in HCC [27]. The eight-gene checkpoint module combines several mechanisms and differs from PD-L1 immunohistochemistry. Identifying which cells express the receptors and ligands, and whether those cells contact one another, would resolve which inhibitory interactions the bulk score reflects.

Regional module measurements can show whether activation, recruitment, and inhibitory expression coexist across sampled regions or occur only at selected sites (Table 3). Adjusted IFN below the reference mean in several regions would identify a candidate activation deficit, a hypothesis for prospective testing in studies of immune stimulation. High IFN together with high checkpoint expression would identify a candidate pattern of activation accompanied by inhibition. Paired pre- and post-treatment biopsies could show whether treatment raises IFN-associated and checkpoint expression together. A single IFN measurement had a higher adjusted ICC than most other modules, though not separable from MHC-I, and cannot establish a tumor-wide activation deficit. For trafficking, cDC1, or myeloid expression, a second region may reveal recruitment-related, dendritic-cell-associated, or inhibitory expression that the first biopsy missed. The memory/IL-7 module describes naive and memory-associated lymphocyte expression within the tumor. The module does not measure the circulating memory-cell reserve or the survival of the sampled cells.

**Table 3.**
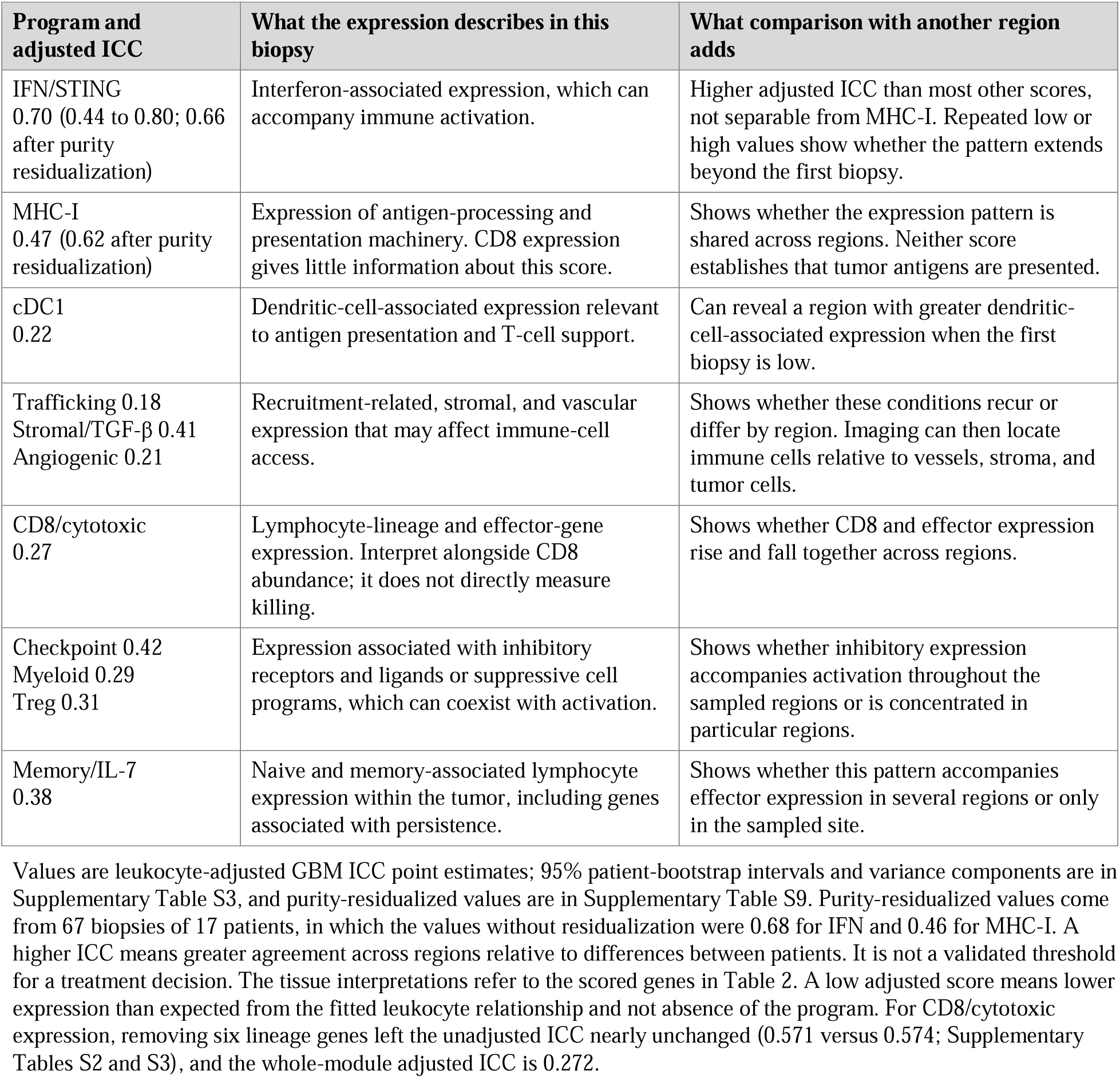
What a GBM biopsy can tell us about selected immune programs.

Measuring supportive and inhibitory programs together preserves distinctions that a single inflamed-versus-uninflamed score would obscure. MHC-I machinery and cDC1 support concern different aspects of antigen presentation, and recruitment, effector transcripts, and cytokine-related support concern different requirements for tumor control. In recurrent GBM, neoadjuvant PD-1 blockade was associated with T-cell and cDC1 activation while suppressive macrophages remained [28], and a pembrolizumab expansion cohort showed T-cell and interferon expression changes without a confirmed survival benefit [29]. These two clinical observations give a biological reason to read the whole panel together and not to rely on a single supportive or inhibitory score.

The patient-level comparison separates a pattern found in every sampled region (G35: cDC1, checkpoint, and Treg expression together) from one that depended on biopsy location (P191: CD8, checkpoint, and stromal expression). Microdissection shows why location matters at a finer scale, because it separates vascular, infiltrative, cellular, and perinecrotic environments that bulk biopsies average [21]. Spatial studies tie myeloid programs to hypoxia and to inflammatory or immunosuppressive niches [13–15]. Our analysis asks how consistently predefined scores characterize one patient across sampled regions. The analysis complements those spatial studies, but a bulk score does not identify the responsible cells or interactions. Multiplex imaging, cell-specific antigen-presentation measurements, and regional functional assays could test whether the combinations in G35 and P191 reflect regulatory-cell interactions, effector localization, or stromal organization.

The higher HCC ICCs indicate greater regional agreement for many immune scores in these cohorts, consistent with the earlier HCC single-region study [17]. Differences in sampling and cohort composition prevent attributing the contrast to cancer type alone. The Treg exception in HCC cohort 1 and the checkpoint estimates in both HCC cohorts argue against applying a cohort-level summary to every program.

Bulk RNA combines cell abundance and cell state, and leukocyte adjustment removes a fitted association without isolating cell-intrinsic function. Without technical replicates, the within-patient variance component cannot separate regional biology from assay variability. Treatment-naive status could not be verified for every GBM sample, and dexamethasone dose and duration were not recorded. The public cohorts analyzed here contain no treatment outcomes and do not evaluate biomarker-guided treatment. The natural killer (NK)/IL-15 and Tfh/IL-21 modules had reduced gene coverage. Only 19 patients contributed to the bulk GBM ICC estimates, seven of them with two biopsies, and six supported disjoint profile recovery. The IFN focus arose after the data were examined, and the order of IFN and MHC-I is unresolved. Several intervals were wide, and the biopsy-number projections assume exchangeable regions, so the analysis does not set a required number of biopsies for a clinical decision.

Regional differences in GBM immune expression mean that a single biopsy can represent the tumor immune microenvironment incompletely. We propose five minimum reporting items for studies that use tissue immune measurements for patient selection or pharmacodynamics (Table 4). The items build on the SITC biomarker consensus [7] and on the sampling-aware checklist proposed for spatial omics in high-grade glioma [19]. Reporting these items would show whether a program behaves as a reproducible patient-level feature or as a regional one before that program guides treatment.

**Table 4.** Proposed minimum reporting for multiregion tissue immune measurements.

| Reporting item | What to report | Finding in this study that motivates the item |
| --- | --- | --- |
| 1. Regions per patient | The number of regions per patient and how regions were chosen (for example, by MRI category). | Profile agreement with other regions improved from 0.28 to 0.32 to 0.42 as one, two, and three biopsies were selected (six GBM patients). |
| 2. Variance components | Between-patient and within-patient variance, and the ICC, on the scale analyzed. | The ICC depends on the variance of each cohort, so values from different cohorts are conditional on their sampling (GBM 0.34, HCC 0.62 and 0.74, GBM-IVY 0.08). |
| 3. Leukocyte adjustment | The ICC before and after adjustment for a named leukocyte composite, and the reliability of that composite. | The median GBM module ICC fell from 0.56 to 0.34 after adjustment, whereas IFN changed from 0.74 to 0.70. The 12-gene composite had a split-half Spearman-Brown reliability of 0.95. |
| 4. Tumor content | Tumor purity or cellularity for each region, and module ICCs before and after residualizing on it. | Residualizing the leukocyte-adjusted scores on WES-inferred purity changed the MHC-I ICC from 0.46 to 0.62 and the IFN ICC from 0.68 to 0.66 (67 biopsies). |
| 5. Paired readouts | CD8, IFN, MHC-I, and checkpoint scores measured in the same regions, with shared interferon-inducible genes identified. | CD8 was nearly uncorrelated with MHC-I ($r = 0.06$), and adjusted IFN and checkpoint scores correlated within patients ($r = 0.60$, and 0.46 without the interferon-inducible genes CD274, IDO1, and PDCD1LG2). |

ICC, intraclass correlation coefficient; WES, whole-exome sequencing. The items are proposals for candidate reporting practice and have not been validated against clinical outcome.

## Methods

### Study design and cohort labels

We performed a retrospective secondary analysis of public tumor transcriptomic datasets with spatially separated samples. GBM denotes the MRI-localized bulk cohort, GBM-IVY the laser-capture anatomical cohort, HCC cohort 1 the Shen cohort, and HCC cohort 2 the Yang cohort. Principal measurements were module scores, regional spread, ICCs, CD8-module correlations, and complete-profile correlations. Exploratory score categories and candidate deficit and inhibitory sets are reported in the Supplementary Methods. No treatment-outcome analysis was performed.

### GBM and GBM-IVY cohorts

The GBM cohort used the batch-adjusted (ComBat-seq) count matrix and sample metadata supplied with Hu and colleagues [20]. Expression values were transformed as log2(x + 1) before gene standardization. The fixed primary cohort included nonnecrotic contrast-enhancing (CE) or nonenhancing (NE) samples from primary, grade IV, isocitrate dehydrogenase (IDH)-wild-type GBM. CE identifies an enhancing MRI region, and NE identifies nonenhancing T2/FLAIR-abnormal infiltrative tissue. The descriptive set contained 79 biopsies from 23 patients. Regional analyses required at least two eligible samples per patient, which left 75 biopsies from 19 patients. Primary disease was not equated with verified treatment-naive status. An erythroid/platelet quality-control sensitivity excluded scores more than two reference SDs above the mean, leaving 72 regions from 19 multiregion patients. The source authors derived tumor purity and gene-level copy-number calls from whole-exome sequencing (WES) with PureCN [30]. Purity and copy number were available for 72 of the 79 biopsies in the descriptive set (22 patients), and 67 of these biopsies from 17 patients had at least two biopsies with a purity estimate.

GBM-IVY used the Ivy Glioblastoma Atlas [21], with normalized fragments per kilobase per million mapped reads (FPKM) transformed as log2(FPKM + 1). The analysis retained 122 canonical anatomical samples from 10 tumors: leading edge, infiltrating tumor, cellular tumor, microvascular proliferation, and pseudopalisading cells around necrosis. It excluded 148 marker-selected or reference-gene samples with a different sampling design. Pseudopalisading samples represent viable perinecrotic tissue and were retained, and their exclusion was examined separately. GBM-IVY was interpreted as an anatomical comparison and not as a replicate of the bulk-biopsy design.

### HCC cohorts

HCC cohort 1 comprised 41 tumor regions from 12 patients in Gene Expression Omnibus accession GSE136711, associated with Shen and colleagues [17]. We converted counts to counts per million (CPM) using all supplied count rows for library size and transformed them as log2(CPM + 1). The downloaded metadata did not provide physical coordinates or region-level necrosis annotations. HCC cohort 2 comprised 75 tumor regions from 14 treatment-naive patients reported by Yang and colleagues [22]. We used the supplied log2(transcripts per million + 1) matrix and excluded 21 normal-liver samples. The source protocol avoided necrotic, fibrotic, hemorrhagic, and cystic areas. The two HCC patient cohorts are independent, and the Yang publication analyzed the Shen cohort separately.

### Module selection and score construction

The fixed gene dictionary contained nine supportive programs, four inhibitory programs, and one checkpoint program (Table 2), chosen to distinguish requirements for completing and sustaining the cancer-immunity cycle [1]. MHC-I describes antigen-processing and presentation machinery, and cDC1 describes dendritic-cell support and cross-presentation. The other modules describe activation, effector support, trafficking, and inhibitory programs. Module names refer to scored gene sets, not cell counts, cytokine concentrations, or direct pathway-function assays. The checkpoint module was included in every continuous profile.

The IFN/STING module contained 22 genes. They encode interferon-stimulated chemokines, signal transducers, interferon regulatory factors, interferon-stimulated effectors, and nucleic-acid sensors, including MB21D1 (CGAS) and TMEM173 (STING). The MHC-I module contained 13 genes for antigen processing and presentation, including PSMB10. Interferon-inducible genes occur in all three modules. Six of them also belong to an interferon-gamma-related expression profile [3]: STAT1 and CXCL9 in IFN/STING, PSMB10 in MHC-I, and CD274, PDCD1LG2, and IDO1 in the checkpoint module. We therefore call the IFN/STING score interferon-associated, without distinguishing type I from type II interferon signaling, and treat the three modules as correlated readouts and not independent ones.

Supplementary Table S1. identifies every dictionary gene and its inclusion. The checkpoint composite averages eight genes; CTLA4, BTLA, and VSIR were unavailable in the MRI reference and were not scored. Comparison cohorts used the MRI-scored sets. The MRI reference contained 165 of 195 audited symbols across biological, leukocyte, and quality-control signatures. NK/IL-15 scored 9 of 16 defined genes and Tfh/IL-21 scored 5 of 9; the other biological modules met the 60% coverage check. Coverage of the MRI-scored sets was complete in the comparison cohorts except for CGAS in HCC cohort 2.

Within each source reference, genes were centered and divided by their population SD, available member genes were averaged with equal weight, and the module average was standardized again. The fixed MRI reference comprised 141 nonnecrotic biopsies, and the primary clinical cohort was a subset. A score of zero denotes the reference mean and not biological inactivity. Cohorts were normalized separately, and absolute scores were not compared between cancer cohorts.

The leukocyte composite used PTPRC, CD52, LCP1, CD53, LAPTM5, CORO1A, ARHGDIB, CD48, CD37, EVL, SASH3, and RAC2 with the same equal-weight standardization. CD3 used CD3D, CD3E, and CD3G, and CD8 used CD8A and CD8B. These are transcriptomic abundance proxies. Each leukocyte-adjusted module score was the standardized residual from regression of its unadjusted score on the leukocyte composite in the source reference. The unadjusted ICC therefore applies to the bulk score, and the adjusted ICC applies to a distinct residual score conditional on this fitted relationship. Adjustment does not identify the cellular source of expression.

### ICC and regional spread

We used a one-way random-effects ICC(1,1) with patient as the grouping factor and an unbalanced method-of-moments estimator. The ICC is the estimated between-patient variance divided by the sum of the between-patient and within-patient variances. It describes the relative reproducibility of a single regional measurement within the sampled cohort, not constancy within every patient. Negative sample estimates were retained. Module intervals used 2,000 patient-bootstrap replicates and abundance intervals used 5,000. Each resampled patient contributed all eligible regions, and repeated draws were treated as distinct clusters. Intervals were conditional on fixed normalization. Reliability bands (below 0.5, poor; 0.5 to 0.75, moderate; 0.75 to 0.9, good; above 0.9, excellent) were applied to the 95% interval and not to the point estimate [24].

Post hoc ICC comparisons used 10,000 paired patient-cluster bootstrap draws (seed 20261003), shared across scores and scales. We compared IFN with each of the 13 other modules on each scale and summarized its bootstrap rank. The unadjusted leukocyte-minus-CD8 contrast was evaluated separately. Supplementary Table S7 reports pointwise and simultaneous intervals, with construction details in the Supplementary Methods. The focus on IFN, and its associated correlations and sensitivity analyses, arose after examining the data and is exploratory. Bootstrap ranks do not remove that selection.

Regional spread was described by the sample SD within each patient. Pooled within-patient variance was the sum of squared deviations from patient means divided by the total within-patient degrees of freedom. It is the within-patient component of the ICC model and does not independently corroborate the ICC. In an unbalanced design, the modeled variance-component sum need not equal the empirical sample variance. The Supplementary Methods and Table S8 define profile RMSE, profile SD, and other descriptive summaries.

Complete-profile correlation measures whether the same modules rank relatively high or low in different regions of one patient. We calculated the Spearman correlation between the 14 module scores for each region pair, took the median within patients, and then took the median across patients. This analysis differs from the CD8-module correlation, which compares two scores across regions, and from the ICC, which compares between-patient and within-patient variance for one score. The Supplementary Methods define additional measures of regional spread and categorical agreement.

### Associations with CD8 and leukocyte expression

Within-patient CD8-module associations ask whether regions with higher CD8 expression than that patient’s average also have higher module expression. Both variables were centered on their patient-specific means. Squared deviations and cross-products were averaged within each patient and then equally across patients. The mean cross-product was divided by the square root of the product of the two mean squared deviations to obtain the correlation. This gives each patient equal total weight and is not an average of individual patient correlations. Figure 2 ranks these correlations within the three module categories, and Supplementary Figure S3 plots selected centered observations with regression slopes computed using the same weights.

The restricted CD8 family covered seven of the 14 modules (IFN/STING, trafficking, checkpoint, angiogenic, stromal/TGF-β, myeloid, and Treg) on the unadjusted and adjusted scales in three bulk cohorts, totaling 42 tests. A dated analysis plan (26 September 2026) fixed the family. The plan followed earlier exploratory GBM work and was not a registered protocol. Two-sided tests used 99,999 patient-level sign flips of cross-products with a plus-one correction, and we calculated Benjamini-Hochberg q values across the 42 tests. Pointwise intervals used 5,000 patient bootstraps. Descriptive unadjusted correlations were also calculated for all 14 modules against CD8 and the leukocyte composite in all three cohorts. These 84 estimates and their 5,000-bootstrap intervals used seed 20261001, without additional discovery testing. Figure 2 and Supplementary Table S4 use this common interval calculation, and Supplementary Table S5 reports the tested family.

To compare supportive and inhibitory programs, we averaged the within-patient CD8 correlations across the eight supportive modules (all supportive modules except CD8/cytotoxic, which shares CD8A and CD8B) and across the four inhibitory modules. The contrast was the supportive mean minus the inhibitory mean. The checkpoint module, a separate category, entered neither mean. Intervals used 5,000 patient bootstraps (seed 20261007), with all 12 correlations recalculated in every resample. This contrast was added after the analysis plan and is exploratory.

Lowest-versus-highest CD8 contrasts used each patient’s sampled extrema, averaging ties before calculating equal-patient mean differences. The cytotoxic sensitivity analysis removed CD3D, CD3E, CD8A, CD8B, CD2, and CD247 and retained GZMA, GZMB, GZMH, PRF1, NKG7, and GNLY. GZMK, CCL5, and IFNG were unavailable in the MRI source. The six-gene measurement can reflect both T-cell and natural killer (NK)-cell expression.

### Sampling depth

Matched sampling selected exactly two, three, or four biopsies per eligible patient in 500 seeded draws for module-profile comparisons, and abundance comparisons used 1,000 two-region draws. These draws check sensitivity to region selection at a fixed biopsy count. The ICC(1,1) remains a single-region estimand at every count. Disjoint profile recovery used the six GBM patients with at least five biopsies. All admissible sets of one, two, or three selected biopsies were compared with exactly two other biopsies, using the mean profile on each side. Split-level measurements were averaged within patient and summarized by the cohort median.

### Anatomical summaries and sensitivity analyses

The paired CE-versus-NE analysis used 11 patients, averaging samples within each MRI category before comparison. Two-sided sign-flip tests enumerated all 2,048 patient sign vectors, and Benjamini-Hochberg correction covered the 14 modules. GBM-IVY anatomical summaries used module scores centered within patients and summarized by structure, and the ordering is descriptive. Module ICCs and profile correlations used all five canonical structures. Complete anatomical outputs are included in the supplementary data.

Sensitivity analyses included erythroid/platelet exclusions, alternative GBM cohort boundaries, tumor purity and copy number, leave-one-patient-out ICC, and leave-one-gene-out scoring. Quality-control subsets did not replace the primary cohort. We also evaluated patients with at least three biopsies, correlation weighting, and regression slopes within and between patients in the 141-biopsy normalization reference. G35 and P191 had the highest and lowest median pairwise profile correlations and the lowest and highest centroid RMSE among all 19 GBM patients on the 14 adjusted modules. These extremes illustrate the range, and all eligible biopsies and modules are shown. Supplementary radar plots display the same values with a radial offset, and polygon area is not analyzed.

### Control analyses

Tumor purity was examined in two ways. We report the biopsy-level Spearman correlation between purity and each adjusted module score in the 72 biopsies. Tests used the slope of an ordinary least squares regression of the score on purity with standard errors clustered on patient, and Benjamini-Hochberg correction across the 14 modules. For the purity-residualized ICC, each adjusted score was regressed on purity by ordinary least squares in the 72 biopsies, and the ICC of the residual was calculated in the 67 multiregion biopsies. Uncertainty came from 2,000 patient-cluster bootstrap resamples of the 22 patients (seed 20261008), with the purity regression refit in every resample. The IFN-minus-MHC-I contrast and the share of resamples in which IFN ranked highest among the 14 modules came from the same resamples. Residualization removes the variance linearly associated with the WES purity estimate and cannot separate tumor-cell-intrinsic expression from composition.

For module size, we drew five genes at random from each module 1,000 times and repeated the full scoring and leukocyte adjustment for every draw (seed 20261006). A module with five scored genes contributed a single set. For the reliability of the leukocyte composite, we split its 12 genes at random into two sets of six, correlated the two half-scores, and applied the Spearman-Brown correction 2r/(1 + r) to the 12-gene length. We evaluated 1,000 splits in the 141-biopsy reference, in the 79 pooled descriptive-set biopsies, and within patients after centering on patient means. To vary adjustment strength, we subtracted k times the fitted leukocyte component before restandardizing (k = 0, 0.5, 0.75, 1, 1.25, and 1.5). A second model used the leukocyte composite and CD3 as covariates. Two-gene controls scored the leukocyte composite from each of its 66 gene pairs and CD3 from each of its three pairs on the unadjusted scale. The checkpoint module was rescored without CD274 and IDO1, and then also without PDCD1LG2.

Copy-number calls for CDKN2A and IFNB1 were the discrete calls supplied by the source authors (−2, homozygous loss; −1, single-copy loss; 0, neutral). A patient was classed as having CDKN2A loss when any biopsy had a call of −1 or −2. Patient-mean adjusted IFN scores were compared between groups with the Mann-Whitney test. To adjust for copy number, we regressed the adjusted IFN score on CDKN2A and IFNB1 copy number in the 67 biopsies with calls and recalculated the ICC of the residual. The glucocorticoid-response score was the standardized mean of FKBP5, TSC22D3, DUSP1, PER1, KLF9, ZBTB16, and SGK1, and a five-gene version omitted DUSP1 and KLF9. We regressed IFN on this score before recalculating its ICC. The score is an unvalidated proxy, and dexamethasone doses were not available.

Region-conditional ICCs removed the CE-versus-NE effect in two ways. The moment-based version subtracted the within-patient estimate of the CE-minus-NE shift and recalculated the one-way ICC. The second version used a random-intercept model with patient as the random effect and region as a fixed effect, fit by restricted maximum likelihood (REML). We wrote the profile likelihood ourselves because a general-purpose mixed-model routine returned boundary solutions with zero between-patient variance for many modules. REML and moment-based estimates agreed (median module ICC 0.35 and 0.34 without the region effect). Region-specific ICCs used patients with at least two biopsies of the same region type. For the projection of regions needed, the Spearman-Brown formula gives the number of regions k whose mean reaches an ICC(1,k) of 0.80 as k = 4(1 − ICC)/ICC, rounded up. Ranges used the limits of the 95% interval. The projection treats regions as exchangeable draws from each patient.

### Statistical reporting conventions

All inferential statements treat the patient as the unit of replication. Regions are repeated measures nested within patients, and n refers to patients unless regions are named. Effect sizes are ICC values, ICC differences, or correlation coefficients, each reported with a 95% percentile interval from the patient-cluster bootstrap unless a permutation q value is stated. Tests for the restricted CD8 family are two-sided sign-flip tests with Benjamini-Hochberg correction. All other contrasts, including the control analyses, are exploratory. Cohort sizes were fixed by the public datasets.

### Reproducibility and data availability

Analyses used the fixed version 1.3 scores. The reproducibility package contains the gene dictionary, cohort labels, figure source values, association families, supplementary analyses, and code. The extensions and control analyses used the same score inputs and ICC estimator. Rescoring from the source count matrix reproduced the archived GBM biopsy scores (maximum absolute difference, 4.6 × 10^−15^). Input checksums, bootstrap settings, and software versions are recorded with the code. Source data are available through the cohort publications and repositories [17,20–22]. [Insert repository name and DOI for the reproducibility package before submission.]

## Declarations

### Ethics

This analysis used publicly available, deidentified data and involved no intervention or interaction with participants and no access to identifiable private information.

### Use of artificial intelligence

Generative AI tools assisted with manuscript drafting, code development, and figure preparation. Numerical results were checked against archived analysis outputs and biopsy-level data. The authors reviewed and take responsibility for the final text, analyses, and interpretations.

### Author contributions

[Insert author contributions using CRediT roles.]

### Data availability statement

All source data are public: Hu and colleagues (GBM), the Ivy Glioblastoma Atlas Project (GBM-IVY), and Gene Expression Omnibus accession GSE136711 and Yang and colleagues (HCC). Derived scores, figure source values, and analysis code are available at [insert repository and DOI].

## Abbreviations

APC: antigen-presenting cell
CD: cluster of differentiation
cDC1: conventional type 1 dendritic cell
CE: contrast-enhancing
CPM: counts per million
FPKM: fragments per kilobase per million mapped reads
GBM: glioblastoma
HCC: hepatocellular carcinoma
ICC: intraclass correlation coefficient
IDH: isocitrate dehydrogenase
IFN: interferon
IL: interleukin
MAD: median absolute deviation
MHC: major histocompatibility complex
MRI: magnetic resonance imaging
NE: nonenhancing
NK: natural killer
PD-1: programmed cell death protein 1
PD-L1: programmed death ligand 1
REML: restricted maximum likelihood
RMSE: root mean square error
SD: standard deviation
SITC: Society for Immunotherapy of Cancer
STING: stimulator of interferon genes
TGF-β: transforming growth factor beta
Th1: type 1 helper T cell
Tfh: T follicular helper cell
Treg: regulatory T cell
WES: whole-exome sequencing.

