## Supplementary Files for "Regional reproducibility of immune expression programs in glioblastoma and implications for single-biopsy assessment"

This file contains Supplementary Methods and Results, Supplementary Figures S1 to S4, and Supplementary Tables S7 to S12. Supplementary Tables S1 to S6 (gene dictionary and coverage, abundance summaries, variance components and ICCs, all-module CD8 and leukocyte correlations, the restricted 42-test family, and all biopsy scores) are supplied as separate data files.

### **Supplementary Methods and Results**

#### ***Additional measures of regional spread and profile recovery***

Unscaled median absolute deviation (MAD) was the median absolute distance from a patient's median biopsy score. For profile MAD, we took the median across the 14 module MADs within each patient and then the median across patients. Adjusted profile MAD was 0.333 in GBM, 0.187 in HCC cohort 1, and 0.226 in HCC cohort 2. Median patient CD8 SDs were 0.554, 0.355, and 0.303, respectively, and the HCC-minus-GBM median-SD intervals included zero. These summaries describe the typical patient, whereas pooled within-patient SD weights patients by within-patient degrees of freedom. The two summaries are different and should not be substituted for each other.

For centroid RMSE, a patient's central profile contained the median score of each module. We calculated the root mean squared distance of each biopsy's 14 scores from that profile and took the median across biopsies. Profile SD was the median sample SD across the 14 adjusted modules. In the separate disjoint-recovery analysis, selected and reference profiles were arithmetic means. At one, two, and three selected biopsies, median profile correlations were 0.282, 0.324, and 0.418 and median RMSEs were 0.855, 0.723, and 0.666. Correlation improved at both steps in all six patients, and at three biopsies patient means ranged from 0.309 to 0.739. All admissible splits were averaged within patient before taking cohort medians.

#### ***Exploratory categories and the restricted correlation family***

Low, intermediate, and high categories used fixed one-third and two-thirds quantiles from the 141-biopsy MRI reference. Low supportive modules and high inhibitory modules formed candidate deficit and inhibitory sets. Checkpoint scores were excluded from these categorical sets but included in all continuous profiles. Jaccard similarity was the number of shared members divided by the number in the union, evaluated for nonempty sets. In disjoint recovery, median similarity was 0.346, 0.342, and 0.356 at one, two, and three selected biopsies. This secondary result was descriptive, and the categories are not validated clinical thresholds.

The GBM CD8 comparisons in the restricted family gave q = 0.055 for IFN and trafficking and q = 0.161 for checkpoint. Low CD8 did not consistently identify a particular adjusted suppressive program (Supplementary Table S5). The expanded all-module estimates are descriptive and do not inherit significance testing from this restricted 42-comparison family.

#### ***Variance components and patient influence***

For unadjusted GBM leukocyte and CD8 scores, estimated between-patient variances were 0.797 and 0.506 and within-patient variances were 0.426 and 0.698, respectively. The modeled variance sums were 1.224 and 1.204, and the empirical total sample variances were 1.179 and 1.176. The difference reflects the unbalanced method-of-moments design. G28 contributed 35.4% of the CD8 within-patient sum of squares, with eight regions, and P191 contributed 13.8%, with four. Such contributions depend on region count. Leave-one-patient-out CD8 ICC ranged from 0.350 to 0.549, compared with 0.620 to 0.682 for leukocytes. The CD8-MHC-I correlation ranged from 0.010 to 0.131.

#### ***Bootstrap comparisons and sensitivity analyses***

For each contrast, the bootstrap SD estimated its standard error. We centered each bootstrap contrast at its observed value, divided by that standard error, and took the largest absolute deviation across the family in each draw. The 95th percentile of these maxima multiplied each standard error to form simultaneous intervals around observed differences. The same procedure was repeated over all 91 pairwise contrasts to account more broadly for choosing IFN after inspecting module rankings. This is a bootstrap maximum standardized-deviation procedure with fixed bootstrap standard errors and not a nested studentized bootstrap.

The 141-biopsy reference comprised 41 patients. For IFN, the pooled, within-patient, and equally weighted between-patient slopes on leukocyte score were 0.665, 0.475, and 0.692. Corresponding values were 0.848, 0.777, and 0.855 for cDC1; 0.914, 0.822, and 0.925 for trafficking; 0.875, 0.829, and 0.899 for myeloid; and 0.753, 0.715, and 0.750 for checkpoint. Replacing each pooled reference slope with its within-patient slope left IFN first in the 75-region set (ICC 0.721), with cDC1, trafficking, and myeloid ICCs of 0.246, 0.230, and 0.295. The original scores remain the primary measurements. Full slope and alternative-ICC outputs accompany the supplementary data.

The correlation-weighting sensitivity analysis pooled patient-centered sums of squares and cross-products, which is equivalent to weighting patient sample covariances by their within-patient degrees of freedom. CD8 correlations changed from 0.063 to 0.079 for MHC-I, 0.456 to 0.421 for IFN, 0.490 to 0.406 for cDC1, 0.475 to 0.424 for checkpoint, 0.480 to 0.444 for Treg, and 0.457 to 0.368 for trafficking. Across all 14 modules, the largest absolute change was 0.100 and some rank positions changed. MHC-I remained weakly associated with CD8. The unadjusted lineage-excluded effector ICC was 0.574, compared with 0.571 for the unadjusted mixed cytotoxic module.

The IFN/STING, MHC-I, and checkpoint modules contained 22, 13, and 8 scored genes, with no pairwise overlap, so their correlations do not arise from identical genes entering both averages. All IFN-MHC-I and IFN-checkpoint correlations used the same equal-patient weighting as the CD8 analyses, and intervals used the paired 10,000-draw patient bootstrap. The adjusted IFN-MHC-I correlation was 0.550 (95% interval, 0.351 to 0.754) within patients and 0.761 across patient means. Within-patient MHC-I associations were 0.424 (0.255 to 0.597) with leukocytes and 0.283 (0.090 to 0.479) with CD3.

Exact CE-NE enumeration reproduced the directional finding: NE checkpoint and trafficking scores were lower in 10 of 11 patients, with Benjamini-Hochberg q values of 0.123 and 0.146. All 14 q values exceeded 0.05.

#### ***Additional results for the control analyses***

In the supportive-versus-inhibitory CD8 contrast, the unadjusted mean correlations were 0.368 across the eight supportive modules and 0.361 across the four inhibitory modules (difference 0.007; 95% patient-bootstrap interval, −0.123 to 0.162). On the adjusted scale the means were 0.115 and 0.190 (difference −0.075; −0.189 to 0.072).

Supplementary Table S9 reports the purity analyses. Table S9 fits the purity regression in all 72 purity-matched biopsies and calculates the ICC in the 67 multiregion biopsies. Residualizing on purity changed the MHC-I ICC by 0.15 (95% interval, −0.06 to 0.32), the myeloid ICC by 0.13 (−0.02 to 0.29), and the IFN ICC by −0.02. Purity correlated positively with MHC-I and negatively with Th1/IL-12 and memory/IL-7, and not with IFN.

Supplementary Table S10A gives the five-gene ICC of every module. Among the 1,000 five-gene IFN subsets, 92.7% exceeded the full-gene MHC-I ICC (0.467), and 96.4% exceeded the highest ICC among the remaining full-gene modules (0.421, checkpoint). On the adjusted scale, IFN had a between-patient variance of 0.848 and a within-patient variance of 0.370, compared with medians of 0.317 and 0.665 across the other 13 modules. IFN therefore combined more between-patient variance with less within-patient variance than the typical module.

Supplementary Table S10C shows that IFN ranked first at every adjustment strength. Its ICC fell from 0.74 to 0.65 as k rose from 0 to 1.5, whereas the median module ICC fell from 0.56 at k = 0 to 0.34 at k = 1 and then rose to 0.40 at k = 1.5. Patient means of the glucocorticoid-response score correlated with the leukocyte composite (rho 0.52; 0.41 for the five-gene score) and not with MHC-I (rho −0.06). When MHC-I and stromal/TGF-β were each residualized on the score, their ICCs stayed at 0.47 and 0.41 (Supplementary Table S10E). The correlation of the score with the leukocyte composite means that the score does not isolate glucocorticoid response.

Supplementary Table S11 reports ICCs conditional on region type. Eleven of the 19 patients had both CE and NE biopsies (40 CE and 35 NE biopsies in all). CE-only ICCs (35 biopsies, 11 patients) and NE-only ICCs (34 biopsies, 13 patients) were 0.75 and 0.78 for IFN and 0.21 and 0.76 for MHC-I. These subsets are small, and we do not interpret the MHC-I difference between them.

Supplementary Table S12 converts each ICC into the number of regions whose mean reaches an ICC(1,k) of 0.80. When the lower limit of the 95% interval was 0.02 or below, the number of regions has no finite upper bound and the range is reported as open-ended.

**Supplementary Figure S1. Radar views of the same patient profiles.**

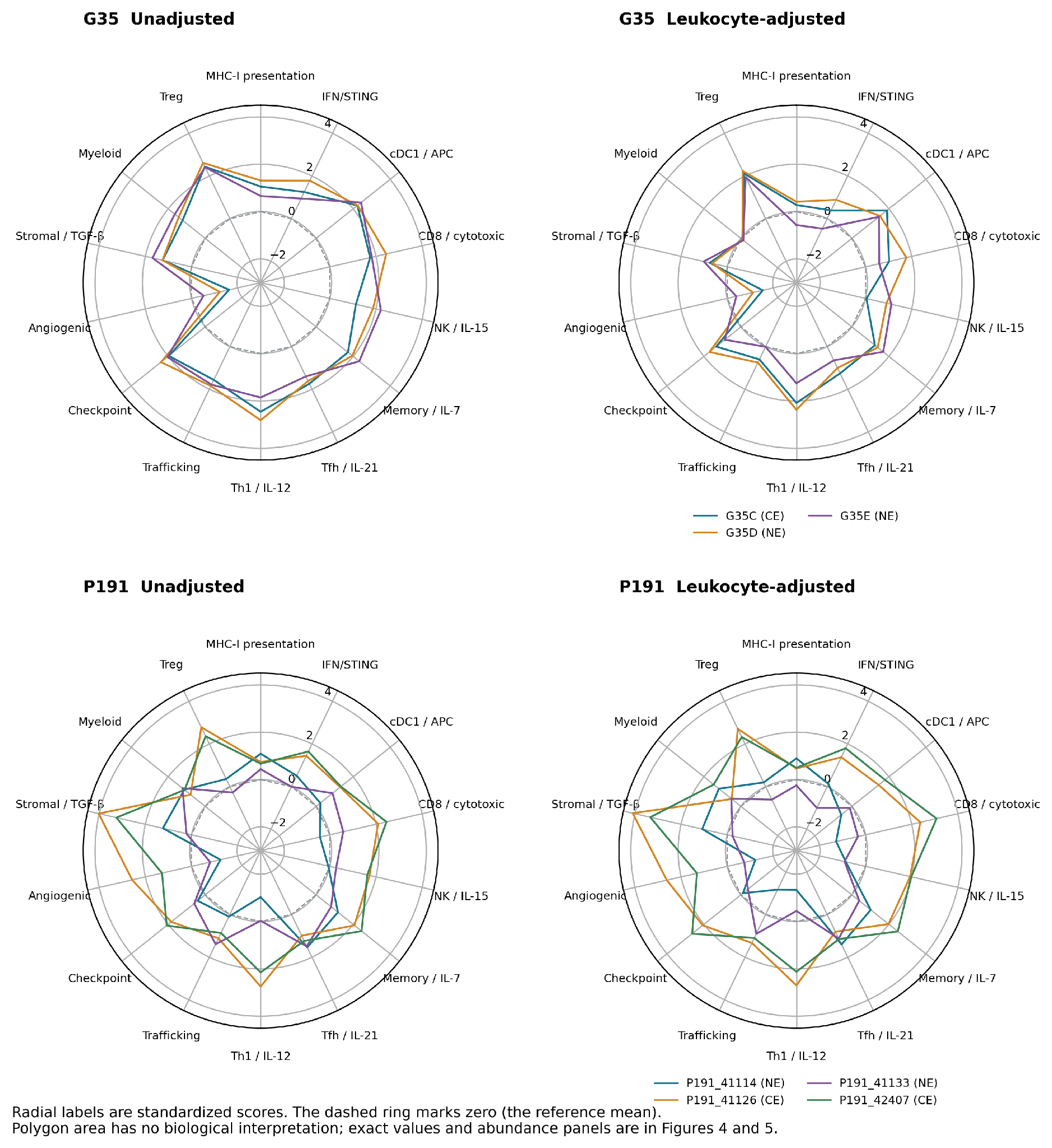

All 14 standardized scores for G35 and P191 are shown before and after leukocyte adjustment. Radial tick labels retain the original score values, and the dashed ring marks zero, the reference mean. The plotting offset allows negative scores on a radial display. Polygon area has no biological interpretation. Abundance measurements and exact linear-scale comparisons are in Figures 4 and 5.

**Supplementary Figure S2. All unadjusted modules compared with both abundance measures.**

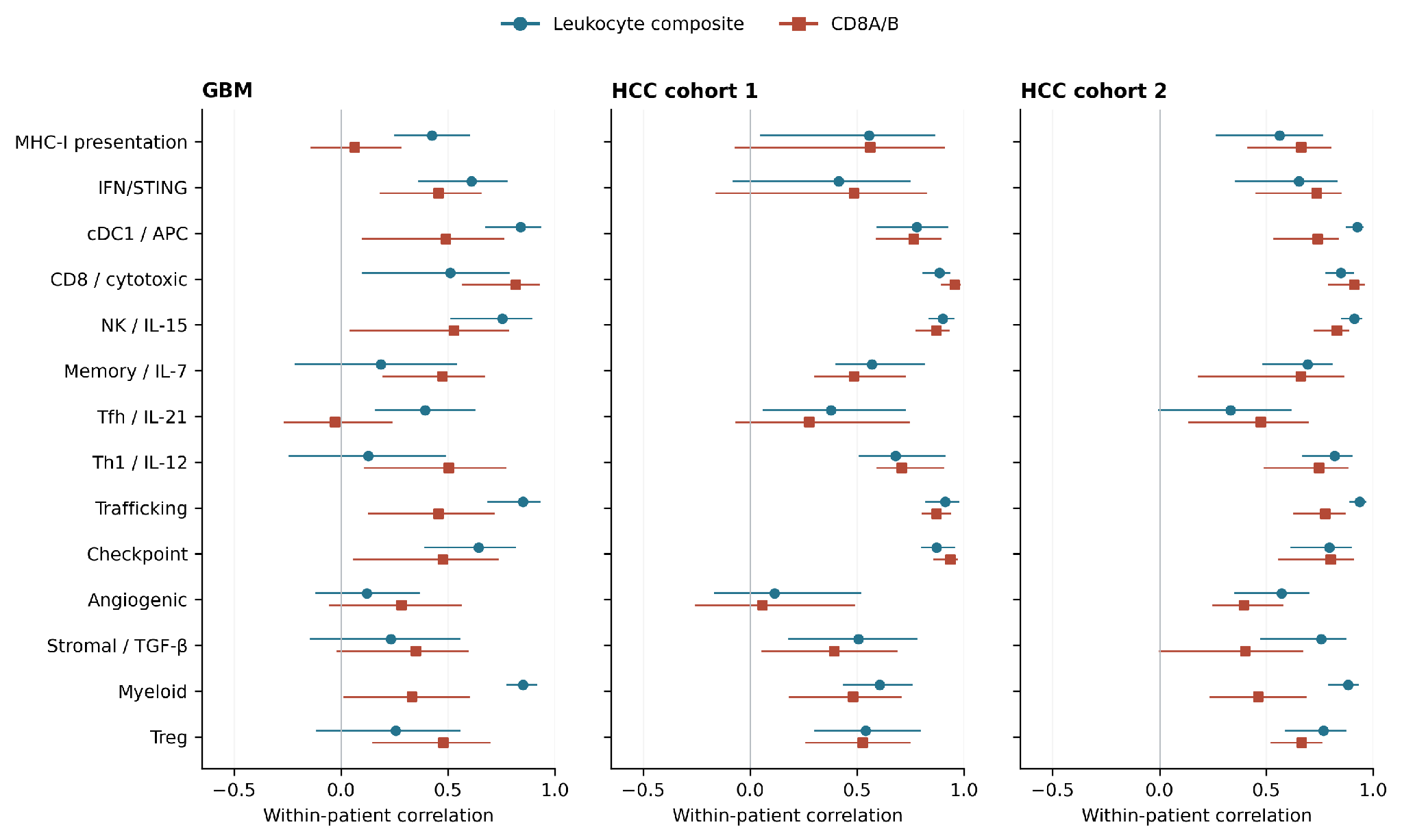

Patient-centered correlations with leukocytes and CD8 for all 14 modules in the three bulk cohorts. Lines are pointwise 95% intervals from 5,000 patient bootstraps with frozen normalization. This expanded display is descriptive and does not extend the restricted 42-test family. CD8A and CD8B contribute directly to the mixed cytotoxic module, so that association includes shared-gene dependence. Exact values are in Supplementary Table S4.

**Supplementary Figure S3. Selected regional module relationships with CD8 and leukocyte expression.**

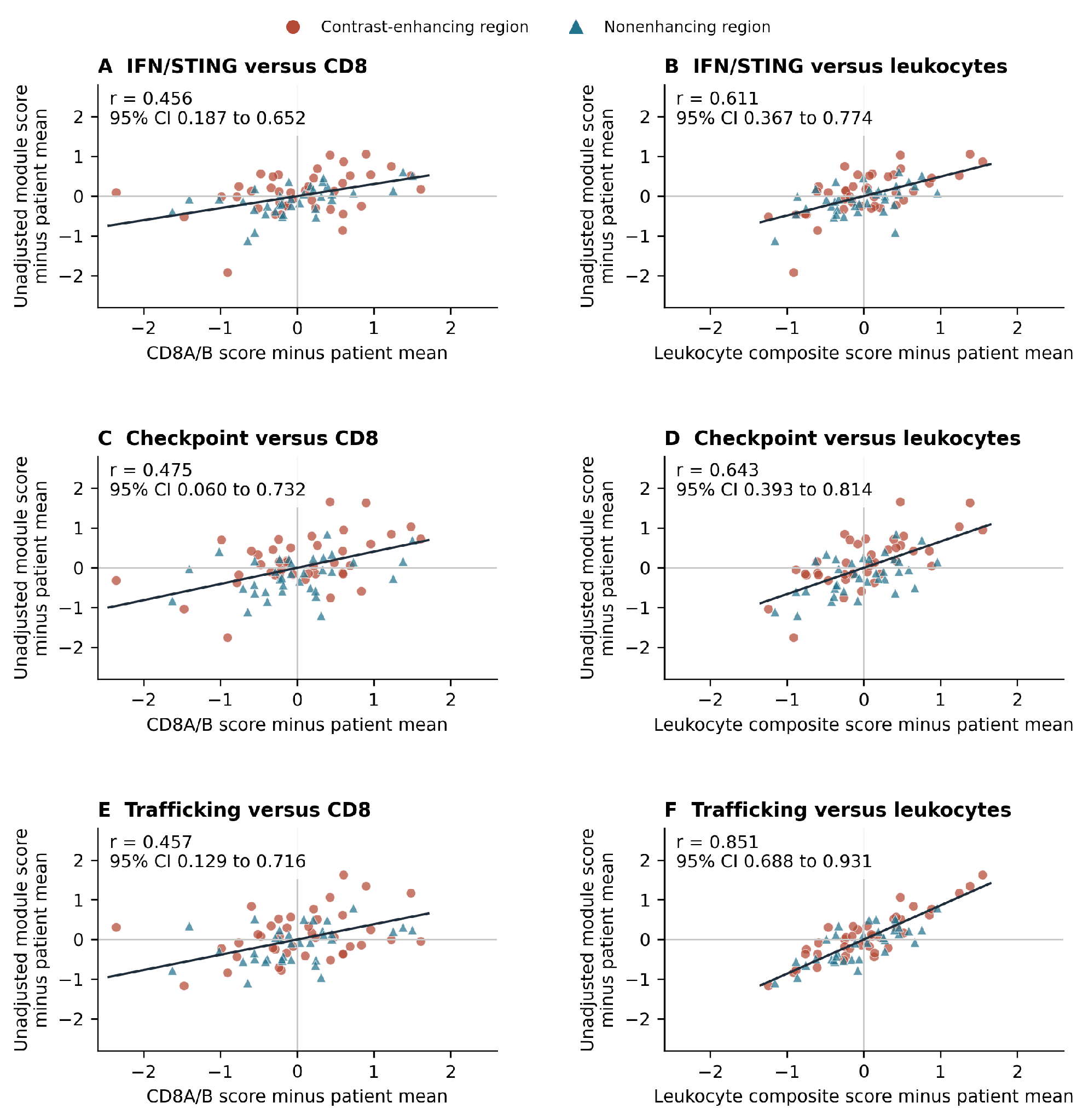

Each point is one of 75 GBM biopsies from 19 patients. IFN/STING, checkpoint, and trafficking scores are plotted against CD8A/B expression in panels A, C, and E and against the 12-gene leukocyte composite in panels B, D, and F. Both coordinates subtract that patient's mean, so each plot shows whether a region above its patient's mean abundance also lies above its patient's mean module expression. The module scores are otherwise unadjusted. Lines use equal patient weights, and r and pointwise 95% bootstrap intervals use the same weighting and the descriptive all-module extension. Symbols distinguish CE and NE biopsies. Positive correlations show regional covariation and not pathway-specific causation. The GBM CD8 q values in the restricted family are 0.055 for IFN, 0.161 for checkpoint, and 0.055 for trafficking (Supplementary Table S5).

**Supplementary Figure S4. Agreement in module ordering and differences in biopsy scores across GBM patients.**

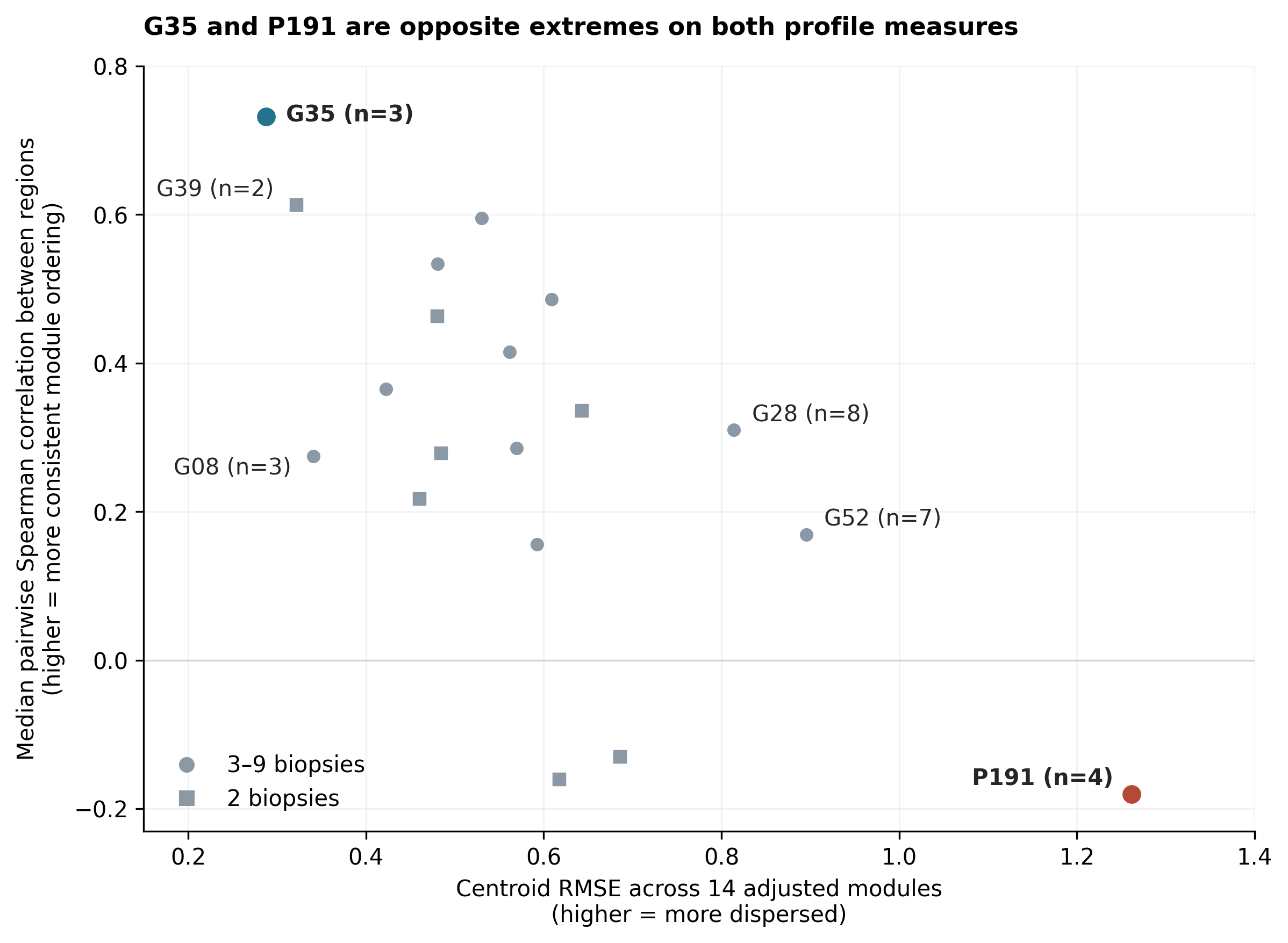

Each point is one GBM patient. The vertical axis gives the median pairwise Spearman correlation across the 14 adjusted module scores, which indicates how consistently the same modules rank high or low across biopsies. The horizontal axis gives the median biopsy distance from the patient's median module profile, calculated as RMSE; larger distances indicate greater regional differences in score values. Circles identify patients with three to nine biopsies, and squares identify patients with two biopsies, who yield only one pairwise correlation. G35 has the highest correlation and smallest distance, and P191 has the lowest correlation and largest distance, among all 19 patients and among the 12 with at least three biopsies. Profile SD is a separate summary of regional spread (Supplementary Table S8). No individual-patient ICC is estimated.

**Supplementary Table S7. Post hoc paired ICC differences and simultaneous intervals**

Panels A and B report IFN-minus-module differences on the adjusted and unadjusted scales in the same 19 patients and 75 regions. Pointwise intervals are 95% bootstrap percentiles. Simultaneous intervals use the maximum standardized bootstrap deviation across all 13 IFN contrasts, separately on each scale. Both use 10,000 paired patient draws and fixed normalization. All 91 pairwise module contrasts were included in a broader selection sensitivity analysis, supplied in the accompanying CSV file; it retained the same three positive adjusted IFN contrasts, with trafficking close to zero. These analyses are separate from the 42-test CD8 family.

**A. Leukocyte-adjusted scores**

| **Comparator** | **Module ICC** | **IFN minus module** | **Pointwise 95%** | **Simultaneous 95%** |
| --- | --- | --- | --- | --- |
| MHC-I | 0.467 | 0.229 | −0.037 to 0.431 | −0.108 to 0.565 |
| cDC1 | 0.220 | 0.476 | 0.073 to 0.792 | −0.055 to 1.008 |
| CD8/cytotoxic | 0.272 | 0.424 | 0.192 to 0.633 | 0.109 to 0.740 |
| NK/IL-15 | 0.369 | 0.326 | 0.042 to 0.583 | −0.068 to 0.721 |
| Memory/IL-7 | 0.379 | 0.317 | −0.106 to 0.704 | −0.285 to 0.919 |
| Tfh/IL-21 | 0.396 | 0.300 | −0.141 to 0.636 | −0.269 to 0.868 |
| Th1/IL-12 | 0.316 | 0.380 | 0.017 to 0.664 | −0.095 to 0.856 |
| Trafficking | 0.178 | 0.518 | 0.197 to 0.790 | 0.088 to 0.949 |
| Checkpoint | 0.421 | 0.275 | −0.000 to 0.527 | −0.105 to 0.654 |
| Angiogenic | 0.207 | 0.489 | 0.240 to 0.686 | 0.159 to 0.819 |
| Stromal/TGF-β | 0.411 | 0.285 | 0.032 to 0.535 | −0.076 to 0.646 |
| Myeloid | 0.294 | 0.402 | 0.027 to 0.685 | −0.076 to 0.881 |
| Treg | 0.312 | 0.384 | 0.043 to 0.726 | −0.121 to 0.889 |

**B. Unadjusted scores**

| **Comparator** | **Module ICC** | **IFN minus module** | **Pointwise 95%** | **Simultaneous 95%** |
| --- | --- | --- | --- | --- |
| MHC-I | 0.558 | 0.178 | −0.003 to 0.314 | −0.054 to 0.409 |
| cDC1 | 0.603 | 0.132 | −0.072 to 0.382 | −0.199 to 0.464 |
| CD8/cytotoxic | 0.571 | 0.164 | −0.027 to 0.377 | −0.133 to 0.461 |
| NK/IL-15 | 0.542 | 0.193 | 0.010 to 0.386 | −0.084 to 0.470 |
| Memory/IL-7 | 0.529 | 0.207 | −0.072 to 0.602 | −0.301 to 0.714 |
| Tfh/IL-21 | 0.591 | 0.144 | −0.062 to 0.377 | −0.178 to 0.467 |
| Th1/IL-12 | 0.447 | 0.288 | 0.038 to 0.586 | −0.124 to 0.700 |
| Trafficking | 0.644 | 0.091 | −0.075 to 0.241 | −0.142 to 0.324 |
| Checkpoint | 0.586 | 0.150 | −0.036 to 0.354 | −0.139 to 0.438 |
| Angiogenic | 0.214 | 0.521 | 0.236 to 0.760 | 0.132 to 0.910 |
| Stromal/TGF-β | 0.448 | 0.287 | 0.051 to 0.599 | −0.122 to 0.697 |
| Myeloid | 0.590 | 0.145 | −0.105 to 0.358 | −0.199 to 0.489 |
| Treg | 0.361 | 0.375 | 0.100 to 0.740 | −0.119 to 0.869 |

The unadjusted leukocyte-minus-CD8 ICC difference was 0.231 (pointwise 95% interval, −0.037 to 0.525). For the adjusted IFN-minus-checkpoint contrast, the unrounded pointwise lower limit was slightly negative, and the rounded value of zero does not establish separation. Simultaneous intervals were not clipped to the parameter space. Bootstrap ranks were descriptive frequencies and not posterior probabilities that a module is best.

**Supplementary Table S8. Profile agreement and regional spread for each GBM patient**

Patients are ordered by profile correlation. SDs use sample denominators, and profile SD is the median SD across the 14 adjusted modules. Profile correlation summarizes all within-patient region pairs, and centroid RMSE summarizes distances from the median profile. Leukocyte and CD8 SDs use unadjusted scores. These shape and distance measures describe different aspects of the same profiles and are not independent biological evidence.

| **Patient** | **Biopsies** | **Profile correlation** | **Centroid RMSE** | **Profile SD** | **Leukocyte SD** | **CD8 SD** |
| --- | --- | --- | --- | --- | --- | --- |
| G35 | 3 | 0.732 | 0.288 | 0.404 | 0.253 | 0.317 |
| G39 | 2 | 0.613 | 0.322 | 0.205 | 0.224 | 0.116 |
| G29 | 4 | 0.596 | 0.530 | 0.532 | 0.362 | 0.211 |
| G77 | 3 | 0.534 | 0.481 | 0.541 | 0.627 | 0.747 |
| G40 | 6 | 0.486 | 0.609 | 0.784 | 0.533 | 0.767 |
| P059 | 2 | 0.464 | 0.480 | 0.426 | 0.594 | 0.554 |
| P232 | 4 | 0.415 | 0.562 | 0.591 | 0.674 | 0.654 |
| P190 | 5 | 0.365 | 0.423 | 0.721 | 0.749 | 0.553 |
| G44 | 2 | 0.336 | 0.643 | 0.637 | 1.250 | 0.297 |
| G28 | 8 | 0.310 | 0.814 | 0.886 | 0.593 | 1.405 |
| G05 | 5 | 0.286 | 0.569 | 0.541 | 0.349 | 0.907 |
| G87 | 2 | 0.279 | 0.484 | 0.445 | 0.090 | 0.288 |
| G08 | 3 | 0.275 | 0.341 | 0.332 | 0.435 | 0.233 |
| G45 | 2 | 0.218 | 0.460 | 0.562 | 0.199 | 0.202 |
| G52 | 7 | 0.169 | 0.895 | 1.008 | 0.818 | 0.570 |
| P129 | 9 | 0.156 | 0.593 | 0.670 | 0.754 | 0.572 |
| P184 | 2 | −0.130 | 0.686 | 0.843 | 0.554 | 0.342 |
| G65 | 2 | −0.160 | 0.618 | 0.842 | 1.759 | 2.091 |
| P191 | 4 | −0.180 | 1.262 | 1.394 | 0.284 | 1.340 |

**Supplementary Table S9. Tumor purity, leukocyte-adjusted module scores, and the IFN/STING versus MHC-I comparison**

Panel A gives the Spearman correlation between WES-inferred purity and each leukocyte-adjusted module score in 72 of the 79 descriptive-set GBM biopsies (22 patients), with Benjamini-Hochberg q values across the 14 modules. It also gives the adjusted ICC in the 67 multiregion biopsies (17 patients) before and after residualizing on purity. Panel B gives patient-bootstrap results from 2,000 resamples of the 22 patients, with the purity regression refit in every resample (seed 20261008).

**A. Association with purity and ICC before and after residualization**

| **Module** | **Spearman rho with purity** | **Benjamini-Hochberg q** | **ICC before** | **ICC after** | **Change** |
| --- | --- | --- | --- | --- | --- |
| IFN/STING | 0.09 | 0.696 | 0.68 | 0.66 | −0.02 |
| MHC-I | 0.40 | 0.012 | 0.46 | 0.62 | 0.15 |
| cDC1/APC | −0.09 | 0.897 | 0.15 | 0.15 | 0.00 |
| Th1/IL-12 | −0.43 | 0.013 | 0.25 | 0.25 | 0.00 |
| CD8/cytotoxic | −0.24 | 0.147 | 0.25 | 0.25 | 0.00 |
| NK/IL-15 | 0.07 | 0.862 | 0.34 | 0.34 | 0.00 |
| Memory/IL-7 | −0.37 | 0.036 | 0.37 | 0.39 | 0.03 |
| Tfh/IL-21 | 0.02 | 0.897 | 0.36 | 0.36 | 0.01 |
| Trafficking/adhesion | 0.17 | 0.279 | 0.13 | 0.15 | 0.02 |
| Adaptive checkpoint | 0.24 | 0.367 | 0.37 | 0.39 | 0.02 |
| Angiogenic | 0.10 | 0.740 | 0.16 | 0.13 | −0.02 |
| Stromal/TGF-β | 0.25 | 0.367 | 0.38 | 0.42 | 0.03 |
| Myeloid | 0.31 | 0.149 | 0.26 | 0.39 | 0.13 |
| Treg | 0.07 | 0.947 | 0.30 | 0.30 | 0.00 |

Purity estimates are from Hu and colleagues and were derived with PureCN. ICC values are on the leukocyte-adjusted scale, and changes were computed from unrounded values. The Benjamini-Hochberg q values apply to patient-clustered regression tests of the purity slope.

**B. Patient-bootstrap comparison of IFN/STING with MHC-I**

| **Quantity** | **Before purity residualization** | **After purity residualization** |
| --- | --- | --- |
| IFN/STING ICC | 0.68 (0.36 to 0.79) | 0.66 (0.30 to 0.78) |
| MHC-I ICC | 0.46 (0.22 to 0.63) | 0.62 (0.31 to 0.75) |
| IFN/STING minus MHC-I | 0.21 (−0.11 to 0.43) | 0.04 (−0.25 to 0.30) |
| Share of resamples with IFN/STING above MHC-I | 0.92 | 0.60 |
| Share of resamples with IFN/STING highest of 14 | 0.80 | 0.55 |
| Change in MHC-I ICC after residualization |  | 0.15 (−0.06 to 0.32) |

Intervals are 2.5th and 97.5th percentiles. Shares are descriptive frequencies from a post hoc comparison and are not probabilities that a module is best.

**Supplementary Table S10. Controls for module size, composite reliability, adjustment strength, checkpoint composition, 9p21 copy number, glucocorticoid response, and abundance gene count**

All values come from the 75 GBM biopsies of 19 patients unless a different set is named. Panel A reduces every module to five randomly drawn genes (1,000 draws, seed 20261006), with leukocyte-adjusted ICCs. Panel B gives split-half reliability of the 12-gene leukocyte composite. Panel C varies the strength of leukocyte adjustment. Panel D removes interferon-inducible genes from the checkpoint module and gives the within-patient correlation of the rescored module with adjusted IFN/STING. Panel E reports 9p21 copy number and a glucocorticoid-response score. Panel F gives two-gene controls for the CD8 comparison on the unadjusted scale.

**A. Module size**

| **Module** | **Genes scored** | **Full-gene ICC** | **Five-gene median ICC (2.5th to 97.5th percentile)** | **Rank** |
| --- | --- | --- | --- | --- |
| IFN/STING | 22 | 0.70 | 0.65 (0.40 to 0.76) | 1 |
| MHC-I | 13 | 0.47 | 0.47 (0.28 to 0.64) | 2 |
| Stromal/TGF-β | 15 | 0.41 | 0.40 (0.25 to 0.58) | 3 |
| Adaptive checkpoint | 8 | 0.42 | 0.40 (0.16 to 0.59) | 4 |
| Tfh/IL-21 | 5 | 0.40 | 0.40 (single set) | 5 |
| Memory/IL-7 | 9 | 0.38 | 0.36 (0.12 to 0.51) | 6 |
| NK/IL-15 | 9 | 0.37 | 0.34 (0.18 to 0.48) | 7 |
| Th1/IL-12 | 5 | 0.32 | 0.32 (single set) | 8 |
| Trafficking/adhesion | 10 | 0.18 | 0.30 (0.06 to 0.54) | 9 |
| Myeloid | 9 | 0.29 | 0.30 (0.03 to 0.48) | 10 |
| Treg | 7 | 0.31 | 0.29 (0.16 to 0.45) | 11 |
| CD8/cytotoxic | 12 | 0.27 | 0.26 (0.15 to 0.39) | 12 |
| cDC1/APC | 8 | 0.22 | 0.24 (0.05 to 0.43) | 13 |
| Angiogenic | 14 | 0.21 | 0.21 (0.08 to 0.36) | 14 |

**B. Reliability of the leukocyte composite**

| **Set** | **Split-half r (median, 2.5th to 97.5th percentile)** | **Spearman-Brown reliability of 12-gene composite** |
| --- | --- | --- |
| 141-biopsy reference | 0.90 (0.82 to 0.94) | 0.95 |
| 79 descriptive-set biopsies, pooled | 0.91 (0.85 to 0.95) | 0.95 |
| Within patient (75 biopsies, patient-centered) | 0.84 (0.76 to 0.90) | 0.91 |

**C. Strength of leukocyte adjustment**

| **Adjustment** | **IFN/STING ICC** | **Next-highest module and ICC** | **Median module ICC** |
| --- | --- | --- | --- |
| 0 times fitted slope (unadjusted) | 0.74 | Trafficking/adhesion 0.64 | 0.56 |
| 0.5 times fitted slope | 0.73 | Trafficking/adhesion 0.54 | 0.47 |
| 0.75 times fitted slope | 0.72 | MHC-I 0.48 | 0.39 |
| 1 times fitted slope (primary) | 0.70 | MHC-I 0.47 | 0.34 |
| 1.25 times fitted slope | 0.67 | MHC-I 0.46 | 0.35 |
| 1.5 times fitted slope | 0.65 | Myeloid 0.48 | 0.40 |
| Leukocyte composite and CD3 as covariates | 0.69 | Stromal/TGF-β 0.43 | 0.29 |

In panel C, k multiplies the fitted leukocyte slope in the 141-biopsy reference before restandardization. The last row uses the leukocyte composite and CD3 as covariates.

**D. Checkpoint module without interferon-inducible genes**

| **Checkpoint gene set** | **Genes** | **Adjusted ICC** | **Unadjusted ICC** | **Within-patient r with adjusted IFN/STING (95% interval)** |
| --- | --- | --- | --- | --- |
| All 8 scored genes | 8 | 0.42 | 0.59 | 0.60 (0.33 to 0.78) |
| Without CD274 and IDO1 | 6 | 0.26 | 0.52 | 0.50 (0.20 to 0.70) |
| Without CD274, IDO1, and PDCD1LG2 | 5 | 0.15 | 0.50 | 0.46 (0.17 to 0.66) |

Within-patient correlations use equal patient weights. The all-gene row repeats the frozen paired-bootstrap interval, and the other two rows use 5,000 patient bootstraps (seed 20261009).

**E. 9p21 copy number and glucocorticoid-response score**

| **Quantity** | **Value** | **Detail** |
| --- | --- | --- |
| CDKN2A loss (any copy loss in at least one biopsy), patients | 19 of 22 |  |
| CDKN2A homozygous loss in at least one biopsy, patients | 15 of 22 |  |
| Patient-mean adjusted IFN/STING, CDKN2A loss versus no loss | 0.34 (n = 19) versus −0.62 (n = 3) | Mann-Whitney p = 0.13; patient-level R-squared 0.14 |
| Patient-mean adjusted IFN/STING, homozygous loss versus other | 0.39 (n = 15) versus −0.18 (n = 7) | p = 0.27; R-squared 0.09 |
| Biopsy-level Spearman rho, CDKN2A copy number versus IFN/STING | −0.22 | p = 0.07 (72 biopsies); calls coded −2, −1, 0, so a negative rho means higher IFN/STING with more copy loss |
| IFN/STING ICC, 67 biopsies with copy number (17 patients) | 0.68 | 0.69 after adjusting for CDKN2A and IFNB1 copy number |
| Glucocorticoid-response score (FKBP5, TSC22D3, DUSP1, PER1, KLF9, ZBTB16, SGK1), unadjusted ICC | 0.44 | 0.50 without DUSP1 and KLF9 |
| Patient-mean glucocorticoid score versus patient-mean IFN/STING (adjusted) | rho 0.18 | p = 0.47 |
| Patient-mean glucocorticoid score versus patient-mean leukocyte composite | rho 0.52 | 0.41 for the five-gene core score |
| IFN/STING ICC after residualizing on the glucocorticoid score | 0.71 (0.70 with the five-gene core score) | Baseline 0.70; MHC-I 0.47 and stromal/TGF-β 0.41 unchanged |

Copy-number calls are the discrete calls supplied by Hu and colleagues. The glucocorticoid-response score is an unvalidated proxy for steroid exposure, and dexamethasone doses were not available.

**F. Two-gene controls for abundance scores**

| **Score** | **Genes** | **ICC (unadjusted)** | **Detail** |
| --- | --- | --- | --- |
| CD8 abundance | CD8A and CD8B (2 genes) | 0.42 | 95% patient-bootstrap interval, 0.10 to 0.68 |
| CD3 abundance | CD3D, CD3E, and CD3G (3 genes) | 0.55 | 0.33 to 0.72 |
| CD3 abundance, two-gene subsets | Each of the 3 gene pairs | 0.51, 0.53, and 0.57 | Same scoring and standardization as the full composite |
| Leukocyte composite | 12 genes | 0.65 | 0.49 to 0.77 |
| Leukocyte composite, two-gene subsets | Each of the 66 gene pairs | Median 0.60 (2.5th to 97.5th percentile, 0.42 to 0.70) | 97% of pairs exceeded the CD8 ICC |

**Supplementary Table S11. ICCs conditional on contrast-enhancing or nonenhancing region type**

All values are leukocyte-adjusted GBM ICCs in 75 biopsies from 19 patients (40 CE and 35 NE). The first column repeats the primary moment-based ICC. The second removes the within-patient CE-minus-NE shift before the moment-based calculation. The third and fourth columns give restricted maximum likelihood (REML) estimates without and with region as a fixed effect. The last two columns use only CE biopsies (35 biopsies, 11 patients) or only NE biopsies (34 biopsies, 13 patients) from patients with at least two biopsies of that type.

| **Module** | **ICC (moments)** | **Region effect removed (moments)** | **ICC (REML)** | **Region fixed, REML** | **CE only** | **NE only** |
| --- | --- | --- | --- | --- | --- | --- |
| IFN/STING | 0.70 | 0.71 | 0.66 | 0.68 | 0.75 | 0.78 |
| MHC-I | 0.47 | 0.49 | 0.48 | 0.49 | 0.21 | 0.76 |
| cDC1/APC | 0.22 | 0.28 | 0.22 | 0.27 | 0.37 | 0.31 |
| Th1/IL-12 | 0.32 | 0.35 | 0.34 | 0.33 | 0.61 | 0.34 |
| CD8/cytotoxic | 0.27 | 0.28 | 0.27 | 0.29 | 0.41 | 0.41 |
| NK/IL-15 | 0.37 | 0.36 | 0.36 | 0.35 | 0.53 | 0.57 |
| Memory/IL-7 | 0.38 | 0.38 | 0.38 | 0.38 | 0.28 | 0.60 |
| Tfh/IL-21 | 0.40 | 0.42 | 0.41 | 0.41 | 0.41 | 0.50 |
| Trafficking/adhesion | 0.18 | 0.24 | 0.19 | 0.24 | 0.33 | 0.04 |
| Adaptive checkpoint | 0.42 | 0.47 | 0.45 | 0.49 | 0.65 | 0.40 |
| Angiogenic | 0.21 | 0.15 | 0.20 | 0.12 | 0.18 | 0.27 |
| Stromal/TGF-β | 0.41 | 0.42 | 0.42 | 0.42 | 0.61 | 0.53 |
| Myeloid | 0.29 | 0.31 | 0.30 | 0.31 | 0.26 | 0.40 |
| Treg | 0.31 | 0.35 | 0.33 | 0.37 | 0.40 | 0.48 |
| Median of 14 | 0.34 | 0.36 | 0.35 | 0.36 | 0.40 | 0.44 |

REML estimates used a profile likelihood implemented for this analysis because a general-purpose mixed-model routine returned boundary solutions with zero between-patient variance for many modules.

**Supplementary Table S12. Regions needed for a mean of regional measurements to reach an ICC(1,k) of 0.80**

The number of regions is k = 4(1 − ICC)/ICC, rounded up, from the Spearman-Brown formula. Point estimates use the GBM ICC on the scale named in each row. Ranges apply the same formula to the 95% patient-bootstrap interval limits of the ICC.

| **Score** | **ICC (point estimate)** | **Biopsies for ICC(1,k) of 0.80** | **Range from 95% interval limits** |
| --- | --- | --- | --- |
| IFN/STING | 0.70 | 2 | 2 to 6 |
| MHC-I | 0.47 | 5 | 3 to 13 |
| Adaptive checkpoint | 0.42 | 6 | 3 to 17 |
| Stromal/TGF-β | 0.41 | 6 | 4 to 18 |
| Median of 14 modules | 0.34 | 8 |  |
| Myeloid | 0.29 | 10 | at least 4 |
| CD8/cytotoxic | 0.27 | 11 | at least 6 |
| cDC1/APC | 0.22 | 15 | at least 4 |
| Angiogenic | 0.21 | 16 | at least 5 |
| Trafficking/adhesion | 0.18 | 19 | at least 7 |
| Leukocyte composite (unadjusted) | 0.65 | 3 | 2 to 5 |
| CD3 abundance (unadjusted) | 0.55 | 4 | 2 to 9 |
| CD8 abundance (unadjusted) | 0.42 | 6 | 2 to 37 |

The projection treats regions as exchangeable draws from each patient. MRI-targeted sampling of CE and NE regions is not exchangeable, so the numbers rank scores by the sampling each needs and do not state a required number of biopsies. Open-ended ranges ("at least") occur where the lower interval limit was 0.02 or below.
